# The Nse5/6-like SIMC1-SLF2 Complex Localizes SMC5/6 to Viral Replication Centers

**DOI:** 10.1101/2022.05.17.492321

**Authors:** Martina Oravcová, Minghua Nie, Nicola Zilio, Shintaro Maeda, Yasaman Jami-Alahmadi, Eros Lazzerini-Denchi, James A. Wohlschlegel, Helle D. Ulrich, Takanori Otomo, Michael N. Boddy

**Affiliations:** Department of Molecular Medicine, The Scripps Research Institute, La Jolla, CA 92037, USA; Institute of Molecular Biology gGmbH (IMB), D-55128 Mainz, Germany; Department of Integrative Structural and Computational Biology, The Scripps Research Institute, La Jolla, CA 92037, USA; Department of Biological Chemistry, David Geffen School of Medicine, University of California, Los Angeles, CA 90095, USA; Laboratory of Genome Integrity, National Cancer Institute, NIH, Bethesda, MD 20892, USA; San Diego Biomedical Research Institute, San Diego, CA 92121, USA

**Keywords:** SIMC1, C5orf25, Nse5, NSMCE5, SMC5, SMC6, polyomavirus.

## Abstract

The human SMC5/6 complex is a conserved guardian of genome stability and an emerging component of antiviral responses. These disparate functions likely require distinct mechanisms of SMC5/6 regulation. In yeast, Smc5/6 is regulated by its Nse5/6 subunits, but such regulatory subunits for human SMC5/6 are poorly defined. Here, we identify a novel SMC5/6 subunit called SIMC1 that contains SUMO interacting motifs (SIMs) and an Nse5-like domain. We isolated SIMC1 from the proteomic environment of SMC5/6 within polyomavirus large T antigen (LT)-induced subnuclear compartments. SIMC1 uses its SIMs and Nse5-like domain to localize SMC5/6 to polyomavirus replication centers (PyVRCs) at SUMO-rich PML nuclear bodies. SIMC1’s Nse5-like domain binds to the putative Nse6 orthologue SLF2 to form an anti-parallel helical dimer resembling the yeast Nse5/6 structure. SIMC1-SLF2 structure-based mutagenesis defines a conserved surface region containing the N-terminus of SIMC1’s helical domain that regulates SMC5/6 localization to PyVRCs. Furthermore, SLF1, which recruits SMC5/6 to DNA lesions, binds SLF2 analogously to SIMC1 and forms a distinct Nse5/6-like complex. Thus, two Nse5/6-like complexes independently regulate human SMC5/6: SIMC1-SLF2 responding to viral challenge and SLF1/2 recognizing DNA damage.

## Introduction

SMC5/6 is the most enigmatic of the three eukaryotic structural maintenance of chromosomes (SMC) complexes, which together maintain genome stability and prevent disease (1–3). All SMC complexes contain heterodimeric SMC ATPases that topologically embrace DNA. With this conserved activity, cohesin (SMC1/3) holds sister chromatids together prior to mitosis, condensin (SMC2/4) compacts chromosomes for mitotic segregation and SMC5/6 resolves DNA recombination intermediates prior to mitosis and meiosis. SMC5/6 recognizes certain DNA structures e.g., supercoiled DNA, and can compact such substrates (4, 5). This function may protect DNA structures from aberrant processing or, present them in a manner that assists their repair (6–10).

In addition to its canonical role in genome stability, human SMC5/6 is emerging uniquely amongst SMC complexes as a critical viral restriction factor. SMC5/6 inhibits the transcription and/or replication of several viruses including hepatitis B (HBV; (11, 12)), herpes simplex (HSV-1; (13)), human papillomavirus (HPV; (14, 15)), Epstein-Barr virus (EBV; (16)), and unintegrated human immunodeficiency virus (HIV-1; (17)). To overcome restriction by SMC5/6, HBV and HIV-1 use the viral proteins HBx or Vpr to target the complex for proteasomal turnover (11, 17). How SMC5/6 combats viruses remains undefined, but it is known that SMC5/6 collaborates with topoisomerases to silence transcription and replication of both HBV and HSV-1 (13). Thus, SMC5/6 may bind supercoiled viral genomes and compact them, making them refractory to access by transcription and replication factors.

The “core” SMC5/6 complex is a well-conserved hexamer, consisting of the SMC5-SMC6 heterodimeric ATPases, the non-SMC-elements 1/3/4 (NSMCE1/3/4 in human and Nse1/3/4 in yeast) heterotrimer that bridges the SMC5/6 headgroups, and the SMC5-associated SUMO ligase NSMCE2 (Nse2 in yeast, reviewed in (2,3,18,19)). In yeast, additional subunits called Nse5 and Nse6 form a heterodimer (Nse5/6) that plays multiple roles in regulating the Smc5/6 core (20–22). Nse5/6 recruits and loads Smc5/6 on chromatin (23–25), and inhibits the ATPase activity of Smc5/6, possibly to stabilize its chromatin association (26, 27). Likely as a result of loading Smc5/6 on DNA and recruiting SUMO, Nse5/6 also “activates” Nse2, promoting its SUMO ligase activity on chromatin (23,24,28,29). Smc5/6 recruitment and activation at DNA damage sites is promoted by an interaction between Nse5/6 and the BRCT domain-containing proteins Brc1/Rtt107, which in turn interact with Rad18 and gamma-H2AX at DNA lesions (18,23,25,30). Consistent with defects in Smc5/6 regulation, cells lacking Nse5/6 are hypersensitive to genotoxins and exhibit aberrant chromosome segregation in mitotic and meiotic cells (8,18,20,23,25,31).

Despite such crucial roles in yeast, Nse5/6 is not evolutionarily conserved at the primary sequence level. In plants, an SMC5/6-associated heterodimer of ASAP1 and SNI1 has been suggested to be the counterpart of Nse5/6 but this was based on its functions and structural modeling (32). In humans, two proteins SMC5/6 localization factor 1 and 2 (SLF1 and SLF2) were proposed to fulfill the roles of Nse5 and Nse6, respectively (33). Consistent with yeast phenotypes, SLF1/2 recruits SMC5/6 to DNA lesions, and its depletion renders cells sensitive to replication stress (33). SLF2 is also required for SMC5/6 localization to unintegrated HIV-1 DNA to induce its transcriptional silencing, which surprisingly does not require SLF1 (17). SLF2 contains an Nse6-like domain in its C-terminus, detectable by algorithms for picking up remote homologies (e.g., HHPred, (34)). But SLF1 bears no sequence similarity to Nse5 (33).

The existing SMC5/6 core and regulatory subunits do not explain its recruitment to and antagonism of viruses. Here, we used proximity labeling and protein purification to define the proteomic environment of SMC5/6 within nuclear foci induced by the large T antigen of polyomavirus SV40. In this context, we identified a poorly characterized protein SUMO interacting motifs containing 1 (SIMC1) as a novel SMC5/6 cofactor. SIMC1 concentrates at SUMO-rich PML nuclear bodies (PML NBs) via its SUMO-interacting motifs (SIMs; (35)) and interacts with SLF2 via its Nse5-like domain, thereby localizing SMC5/6 to polyomavirus replication centers (PyVRCs). Structural analyses combining AlphaFold prediction and cryo-electron microscopy (cryo-EM) reveal the Nse5/6- like structure of the SIMC1-SLF2 complex. In addition, structure-based mutational analysis establishes the importance of SIMC1’s Nse5-like domain in SMC5/6 regulation. Overall, we find that SIMC1 and SLF1 form distinct Nse5/6- like complexes with SLF2 to direct the antiviral or DNA repair responses of human SMC5/6, respectively.

## Results

### SIMC1 identified proximal to SMC5 within LT-induced subnuclear foci

The large T antigen (LT) of SV40 induces nuclear foci in cells that contain DNA replication and repair factors related to the viral lifecycle (36). Moreover, although functionally unexplored, a previous proteomic analysis of LT-associated proteins identified several subunits of SMC5/6 (37). Consistent with this, we detected SMC5/6 in nuclear foci that colocalized with SV40 LT in HEK293T cells (**Figure 1a**). HEK293 cells lack SMC5/6 foci but expression of LT led to the emergence of puncta containing both SMC5/6 and LT. Thus, SMC5/6 may be a new cellular factor involved in the SV40 lifecycle.

**Figure 1:**
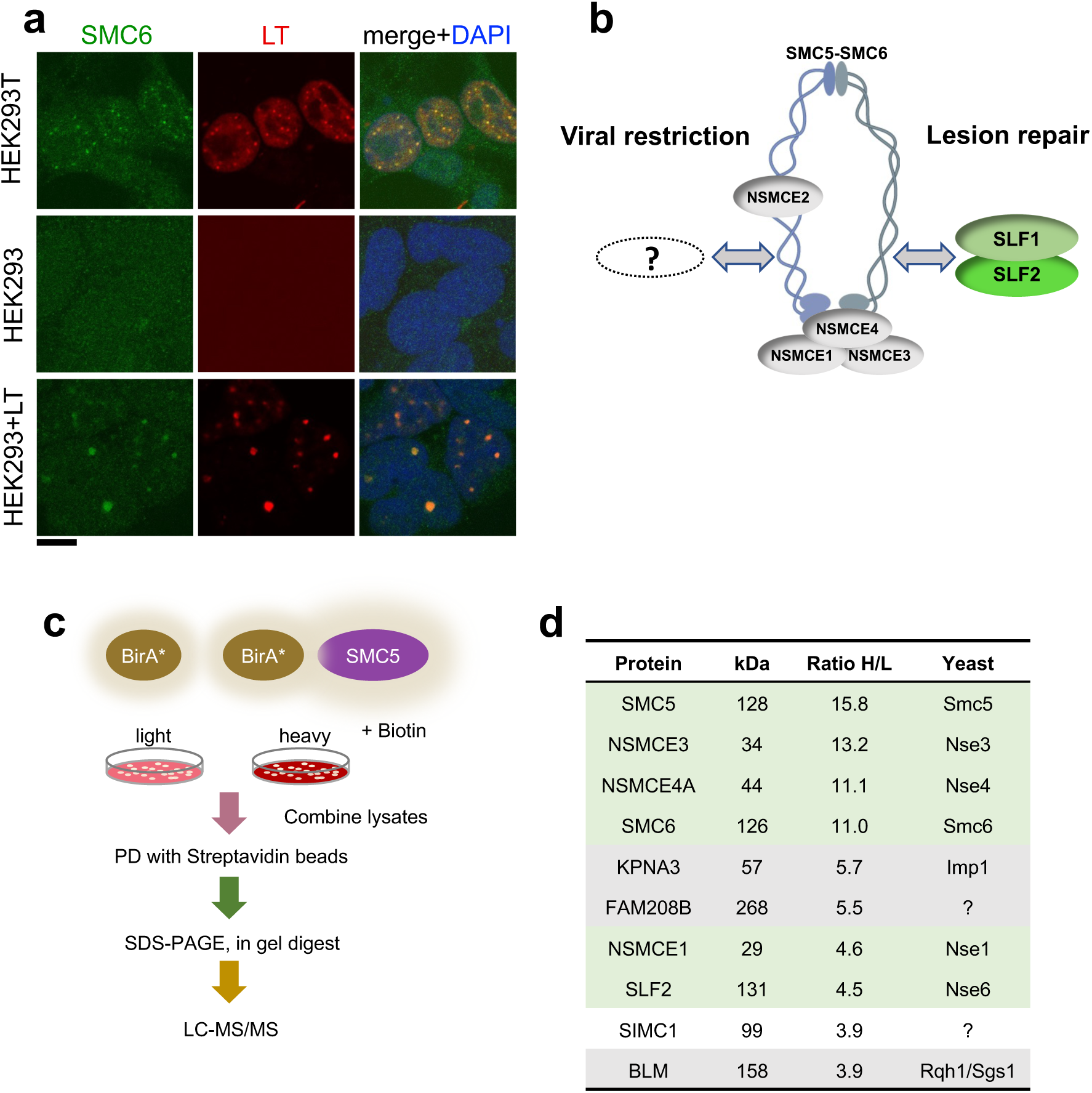
SIMC1 detected in proximity of SMC5 in LT-induced foci. **(a)** Representative immunofluorescence images of HEK293T, HEK293 cell lines and HEK293 cells transiently expressing SV40 large T (LT) antigen. Cells were fixed, stained with SMC6 (green), LT (red) antibodies and DAPI (blue). Scale bar 10 μm. **(b)** Schematic of SMC5/6 complex subunits showing the SLF1/SLF2 cofactors recruiting the complex to DNA lesions and depicting the hypothesis of SMC5/6 cofactor involved in SV40 virus lifecycle. **(c)** Experimental design for human SMC5 BioID with SILAC. **(d)** The top SMC5 interacting proteins identified in BioID screen are listed with their ratio of heavy (H) to light (L) enrichment. Where known, yeast orthologues are listed. The green rows contain known SMC5/6 subunits, grey ones are possible novel interactors of human SMC5. BioID datasets are available in the PRIDE database (https://www.ebi.ac.uk/pride/archive) under the accession code PRIDE PXD033923 and in Supplementary file 1.

Based on the foregoing, we reasoned that virus-related SMC5/6 cofactors may be concentrated within LT-induced nuclear foci (**Figure 1b**). Therefore, we used proximity labeling with biotin and SILAC-based mass spectrometry to identify proteins in the environment of human SMC5 that was fused to the promiscuous biotin ligase BirA* and expressed in HEK293T cells ((38), **Figure 1c**). We defined the top-ranking hits by the SILAC ratios of proteins recovered in SMC5-BirA* versus the NLS-BirA* control purifications (**Figure 1d**, Supplementary file 1 and PRIDE database: PXD033923). Proteins on the list provide high confidence in the data, as 5 of the top 9 most enriched SMC5 interactors are known components of the SMC5/6 core complex (**Figure 1d**). In addition, the presumptive Nse6 orthologue SLF2 was detected but the suggested Nse5 counterpart SLF1 was missing. Instead, a poorly characterized protein called SIMC1 (C5orf25) was detected with similar enrichment as SLF2 (**Figure 1d**).

### SIMC1 contains a yeast Nse5-like domain

SIMC1 is a modular protein containing tandem SIMs within an N-terminal large intrinsically disordered region (IDR) and a predicted alpha-helical rich C-terminal region (**Figure 2a**). Standard sequence searches failed to identify homologous proteins in lower eukaryotes such as yeast. Therefore, we conducted remote homology searches on the HHpred server (34). Searching against yeast and *A. thaliana* databases returned yeast Nse5 and plant SNI1 as the most probable hits in each species for SIMC1 C-terminal amino acids 467-802 (**Figure 2 – figure supplement 1**). Consistent with previous reports (33) a search with the putative Nse5 orthologue SLF1 did not identify Nse5. These results led us to hypothesize that SIMC1 is the Nse5-like regulator of human SMC5/6 that directs SMC5/6 to antagonize viral infections.

**Figure 2:**
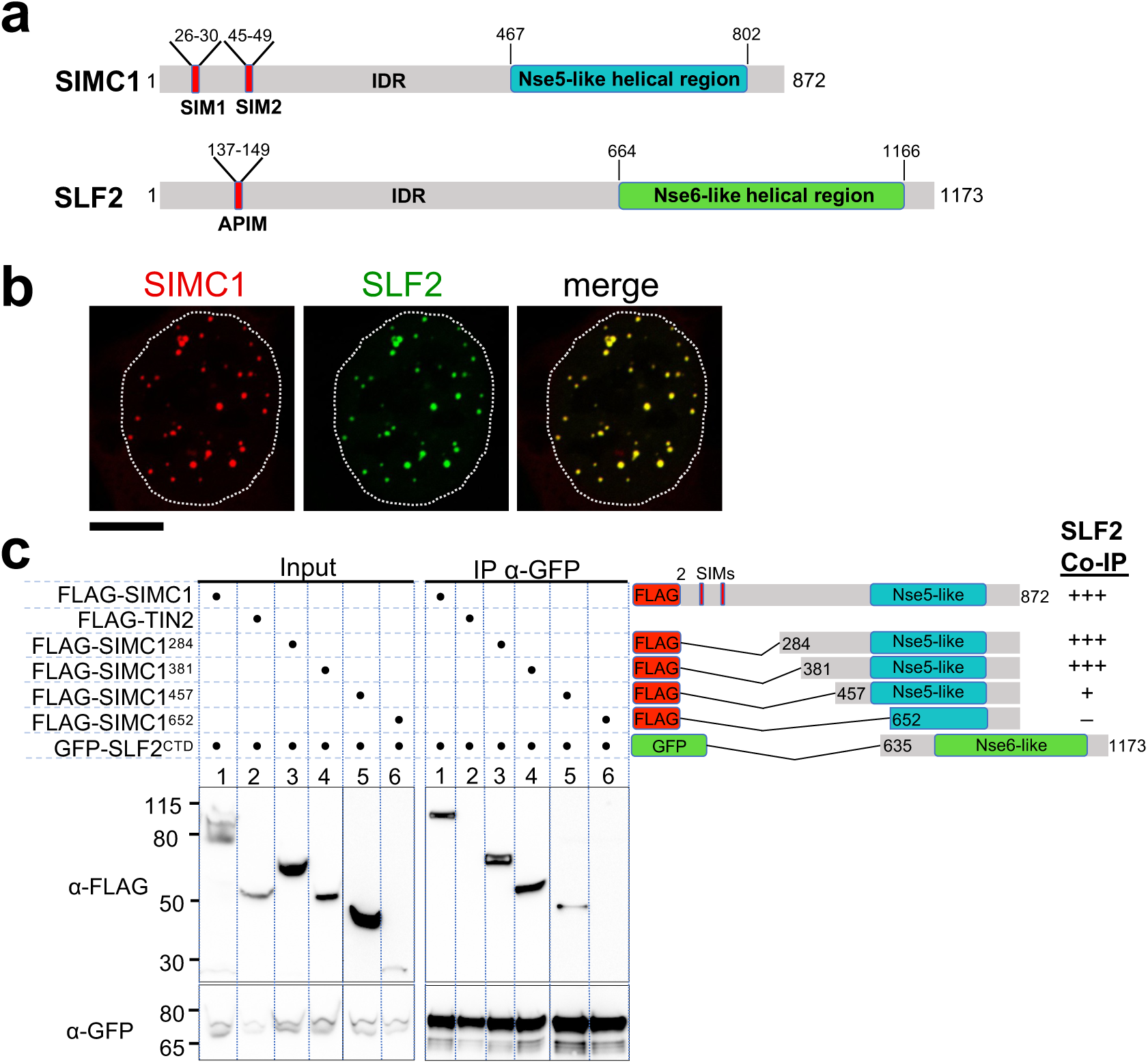
The Nse5- and Nse6-like domains of SIMC1 and SLF2 interact. **(a)** Schematic representation of SIMC1 and SLF2. SIM1/2, SUMO interaction motifs; IDR, intrinsically disordered regions; APIM, AlkB homologue 2 PCNA-interacting motif. **(b)** Representative immunofluorescence images of U2OS cells overexpressing mCherry-SIMC1 and GFP-SLF2 that were fixed and stained with DAPI. Dotted outline marks nucleus. Scale bar 10 µm. **(c)** Western blot of GFP-Trap immunoprecipitation from HEK293 cells transfected with plasmids expressing the indicated combinations of proteins. Truncation constructs of SIMC1 and SLF2 are represented schematically including the status of SIMC1-SLF2 interaction i.e., *** > *, – undetectable. Full and unedited blots provided in Figure 2-source data 1.

### SIMC1 and SLF2 interact via their Nse5- and Nse6-like regions

In all species tested, Nse5/6-like complexes are composed of two alpha-helical domain-containing proteins that form an obligate heterodimer. We, therefore, investigated if and how SIMC1 and SLF2 interact. When co-expressed in cells, full-length SIMC1 and SLF2 colocalize in subnuclear foci and specifically co-immunoprecipitate (**Figure 2b, Figure 2 – figure supplement 2**). Co-expressed SIMC1 and SLF2 are more abundant than when each is expressed with controls, which likely reflects the stabilization of each protein when in complex (**Figure 2 – figure supplement 2**).

Next, we sought to determine the interacting regions of SIMC1 and SLF2. For SLF2, we tested a C-terminal construct (SLF2^CTD^; 635-1173) that contains the Nse6-like region previously reported to interact with SLF1 (39). This construct supports binding to full-length SIMC1 (**Figure 2c**). For SIMC1, we tested four N-terminal truncations, starting with residues 284, 381, 457 or 652. The results showed that SIMC1 constructs 284 and 381 stably interact with SLF2^CTD^ (**Figure 2c**). However, the shortest construct that lacks the N-terminal part of the SIMC1 Nse5-like region (652) was poorly expressed (**Figure 2c**). Whilst SIMC1 construct 457 interacts with SLF2^CTD^ and contains the entire Nse5-like region defined by HHpred, the binding appears weakened (**Figure 2c**). Thus, the minimum SLF2-interacting region of SIMC1 with full binding capacity is located between residues 381 and 872. In parallel, we succeeded in co-purifying an apparently heterodimeric complex of SIMC1^284^ and SLF2^CTD^ proteins expressed in insect cells (**Figure 2 – figure supplement 3**). Thus, the C-terminal domains of SIMC1 and SLF2 directly interact to form a stable complex, supporting our hypothesis that the SIMC1-SLF2 complex is an Nse5/6-like regulatory factor for human SMC5/6.

### SIMC1 interacts with SMC5/6 and SUMO pathway factors

To probe the local environment where SIMC1 functions, we used proximity labeling and mass spectrometry in cells stably expressing either Myc-BirA* or Myc-BirA*-SIMC1 together with SLF2. Because the biotinylation radius of BirA* is restricted to ∼10 nm, direct protein contacts within multiprotein complexes can be detected in native but not denaturing conditions (40, 41). Indeed, in native BioID conditions, SLF2 and all known subunits of the SMC5/6 complex were detected (**Figure 3a**); whereas under the denaturing conditions of BioID, SLF2 but no other SMC5/6 complex subunits were recovered (**Figure 3b**). These results confirm our SMC5 BioID results (**Figure 1d**), and together, they establish SIMC1 as an SMC5/6 subunit.

**Figure 3:**
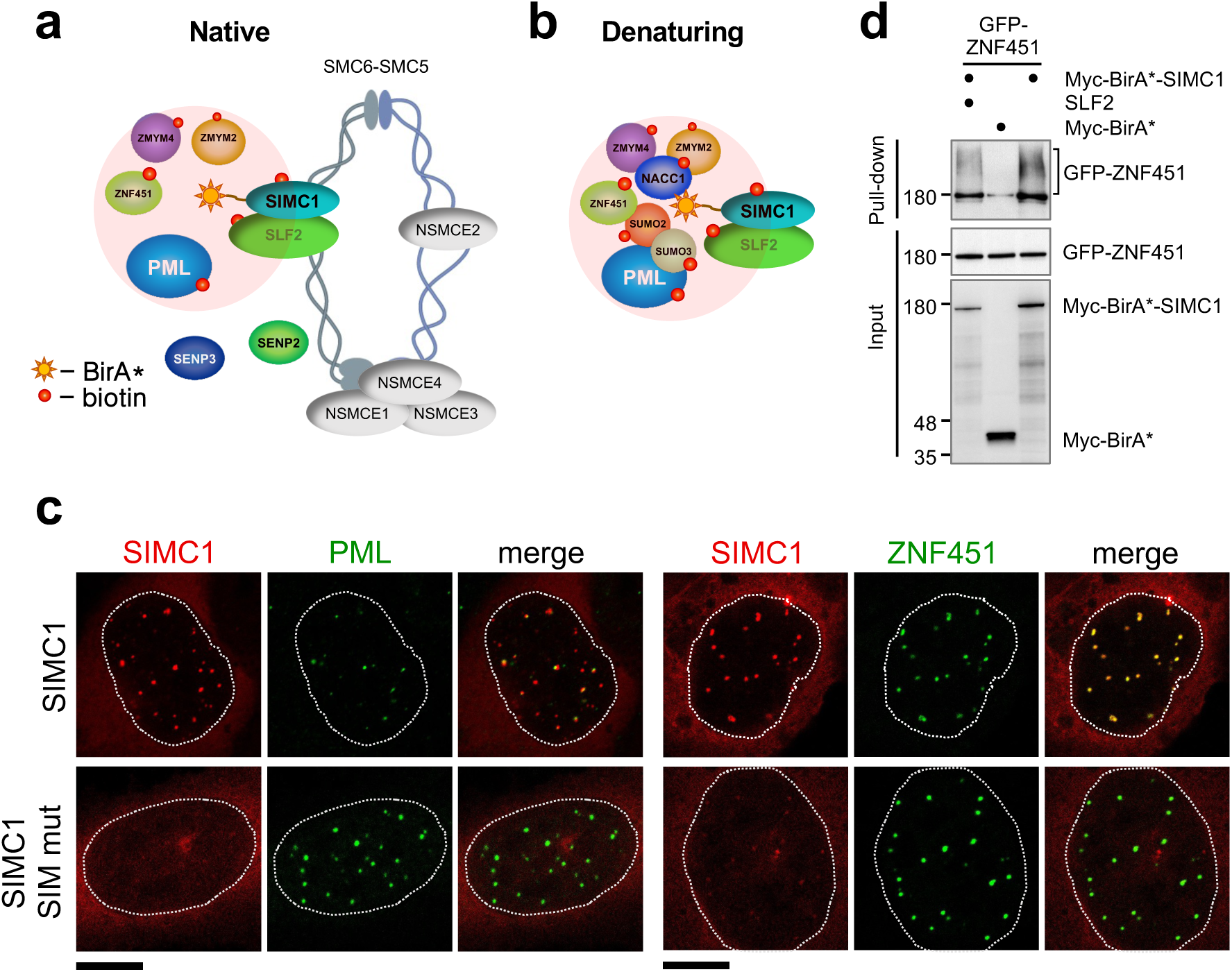
SIMC1 interacts with SMC5/6 and SUMO pathway factors. **(a, b)** A subset of SIMC1 interacting proteins identified by BioID screen carried out under native (a) or denaturing (b) lysis conditions, respectively. Pink circle illustrates the biotinylation radius of BirA*. BioID datasets are available in the PRIDE database (https://www.ebi.ac.uk/pride/archive) under the accession code PRIDE PXD033923 and in Supplementary file 2. **(c)** Representative immunofluorescence images of U2OS cells overexpressing mCherry-SIMC1 or mCherry-SIMC1 SIM mut (mutations: FIDL to AADA (26-30 aa) and VIDL to AADA (45-49 aa)) that were stained for PML (left three panels); or were co-expressing GFP-ZNF451 (right three panels). The cells were stained with DAPI to mark nucleus (dotted lines). Scale bar 10 µm. **(d)** HEK293 cells overexpressing ZNF451, BirA*-SIMC1, SLF2, BirA* control in indicated combination were cultured in the presence of 50 μM supplemental biotin. The total cell extracts and streptavidin pulldown were analyzed by western blot using anti-GFP or Myc antibodies. Full and unedited blots provided in Figure 3-source data 1.

In both conditions, multiple SUMO pathway factors were also identified as SIMC1 proximal, including PML, SUMO2/3 and ZNF451 (**Figure 3a, b,** Supplementary file 2 and PRIDE database: PXD033923), a SUMO2/3 specific E3 ligase that can drive SUMO2/3 chain formation (42, 43). Supporting this, SIMC1 was previously identified using a computational string search for proteins that contain multiple SIMs, and was found to bind SUMO2 polymers *in vitro* via its tandem N-terminal SIMs (35, 44).

We further tested SIMC1’s interaction with the BioID hits PML and ZNF451. Expressed SIMC1 forms nuclear foci that colocalize with both endogenous PML NBs and expressed ZNF451, in a manner dependent on the SIM motifs of SIMC1 (**Figure 3c**). Moreover, epitope-tagged ZNF451 co-purifies with Myc-BirA*-SIMC1 but not a Myc-BirA* control (**Figure 3d**). The SIMC1 interaction with ZNF451 is not enhanced by co-expression with SLF2, consistent with SIMC1 having direct SIM-mediated contacts with both SUMO and SUMO pathway factors (**Figure 3d**). These data validate the interaction of SIMC1 with the SUMO pathway and link SMC5/6 and SUMO-regulated PML NBs via SIMC1.

### SIMC1 recruits SMC5/6 to SV40 replication centers

Because SV40 has been reported to localize to PML NBs (45), SIMC1 may recruit SMC5/6 to SV40 viral replication centers. To test this, we first confirmed the colocalization of SMC5/6, PML, and SV40 LT in permissive HEK293 cells transfected with replication competent SV40. As anticipated, SV40 replication centers (as marked by LT) contain endogenous SMC5/6 and often colocalize with PML NBs (**Figure 4a**). Thus, SMC5/6 is a novel DNA repair and replication factor found at sites of SV40 replication. We then generated SIMC1 knockout HEK293 cells, which are viable and show no overt changes to cell cycle distribution or signs of spontaneous DNA damage, as determined by FACS analysis and comet assay, respectively (**Figure 4 – figure supplement 1**).

**Figure 4:**
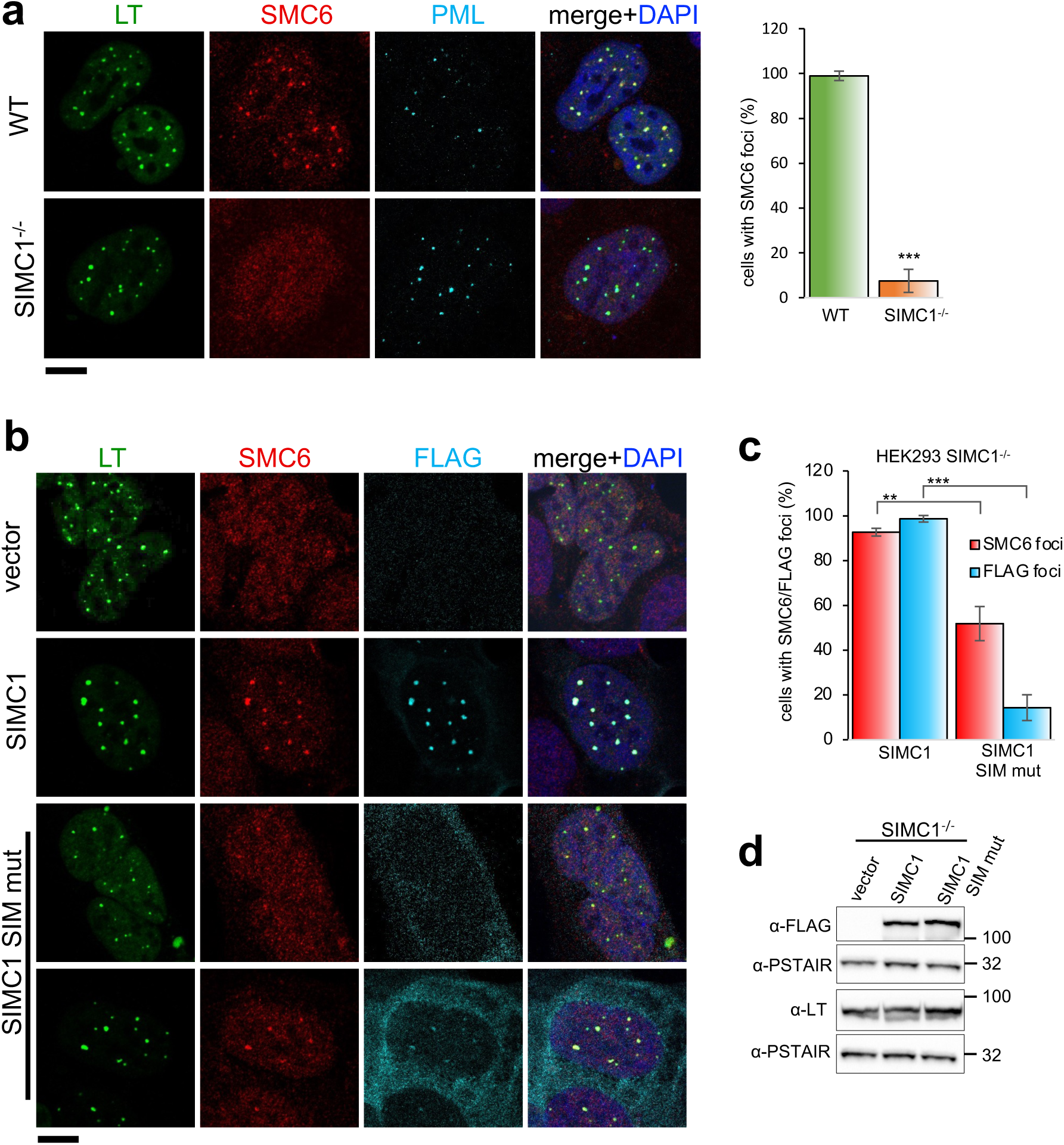
SIMC1 recruits SMC5/6 to SV40 replication centers. **(a)** Representative immunofluorescence images of SV40 LT (green), SMC6 (red) and PML (light blue) in HEK293 and HEK293 SIMC1^-/-^ cells fixed 48 h after SV40 transfection. Scale bar 10 μm. The bar graph shows relative quantification of the number of cells containing SMC6 foci. A minimum of 175 cells with at least four SV40 LT foci were counted for each cell line. Primary data for graph in panel **a** provided in Figure 4-source data 1. **(b)** Representative immunofluorescence images of HEK293 SIMC1^-/-^ cells with integrated empty vector or vectors expressing FLAG-SIMC1 or FLAG-SIMC1 SIM mut, respectively, 48h after SV40 transfection. SV40 LT (green), SMC6 (red), FLAG (light blue); Scale bar 10 μm. **(c)** Relative quantification of the number of cells containing SMC6 and FLAG foci (representative images shown in panel **b**). A minimum of 225 cells with at least four SV40 LT foci were counted for each cell line. Primary data provided in Figure 4-source data 2. **(d)** Immunoblot from HEK293 SIMC1^-/-^ cells with integrated empty vector or vector expressing FLAG-SIMC1 or SIMC1 SIM mut, respectively, and transfected with a plasmid carrying SV40 genome. Cells were harvested 48 h after transfection. PSTAIR serves as a loading control. Full and unedited blots provided in Figure 4-source data 3. Statistics in **(a, c)**: Means and error bars (s.d.) were derived from a minimum of 3 independent SV40 transfections representing biological replicates. (∗) P < 0.05; (∗∗) P < 0.005; (∗∗∗) P < 0.0005; (n.s.) P > 0.05 (two-tailed unpaired t-test).

In SIMC1 null cells, SMC5/6 localization to SV40 replication centers is abolished (**Figure 4a**). Re-expression of SIMC1 restored SMC5/6 colocalization with SV40 LT and SIMC1 (**Figure 4b, c**). Despite similar expression levels as wild-type, SIMC1 with mutated SIMs only partially restored SMC5/6 colocalization with SV40 LT, indicating that it is hypomorphic (**Figure 4b-d**). In addition, the SIM mutant of SIMC1 failed to colocalize with PML and ZNF451 NBs (**Figure 3c**). Thus, SIMC1 and its interaction with SUMO are required for SMC5/6 localization to PyVRCs.

### SIMC1 and SLF2 form an Nse5/6-like structure

To gain insights into the molecular mechanism underlying SMC5/6 regulation, we analyzed the structures of the Nse5- and Nse6-like domains of SIMC1 and SLF2. AlphaFold structure prediction has provided highly reliable structures for proteomes (46). The AlphaFold model of SIMC1 confirms that the N-terminus is disordered and reveals that the Nse5-like region forms an α-solenoid-like helical structure. *S. cerevisiae* Nse5 (ScNse5) also has an α-solenoid-like structure (27, 29); therefore, we compared SIMC1 and ScNse5 structures. Because automated superimposition did not work well, we used remote sequence homology. The HHpred matches are limited to the core region of SIMC1 ranging from α5 to α14 (**Figure 5 – figure supplement 1**). Mapping the matched α-helices on the structures indicated that the matched regions are topologically similar (**Figure 5a**). Manual alignment of the matched regions resulted in superimposing the matched region α5-α14 and α1-α4 as well. Thus, SIMC1 is mostly similar to Nse5 except for the C-terminus (α15-α17), which is absent in ScNse5.

**Figure 5.**
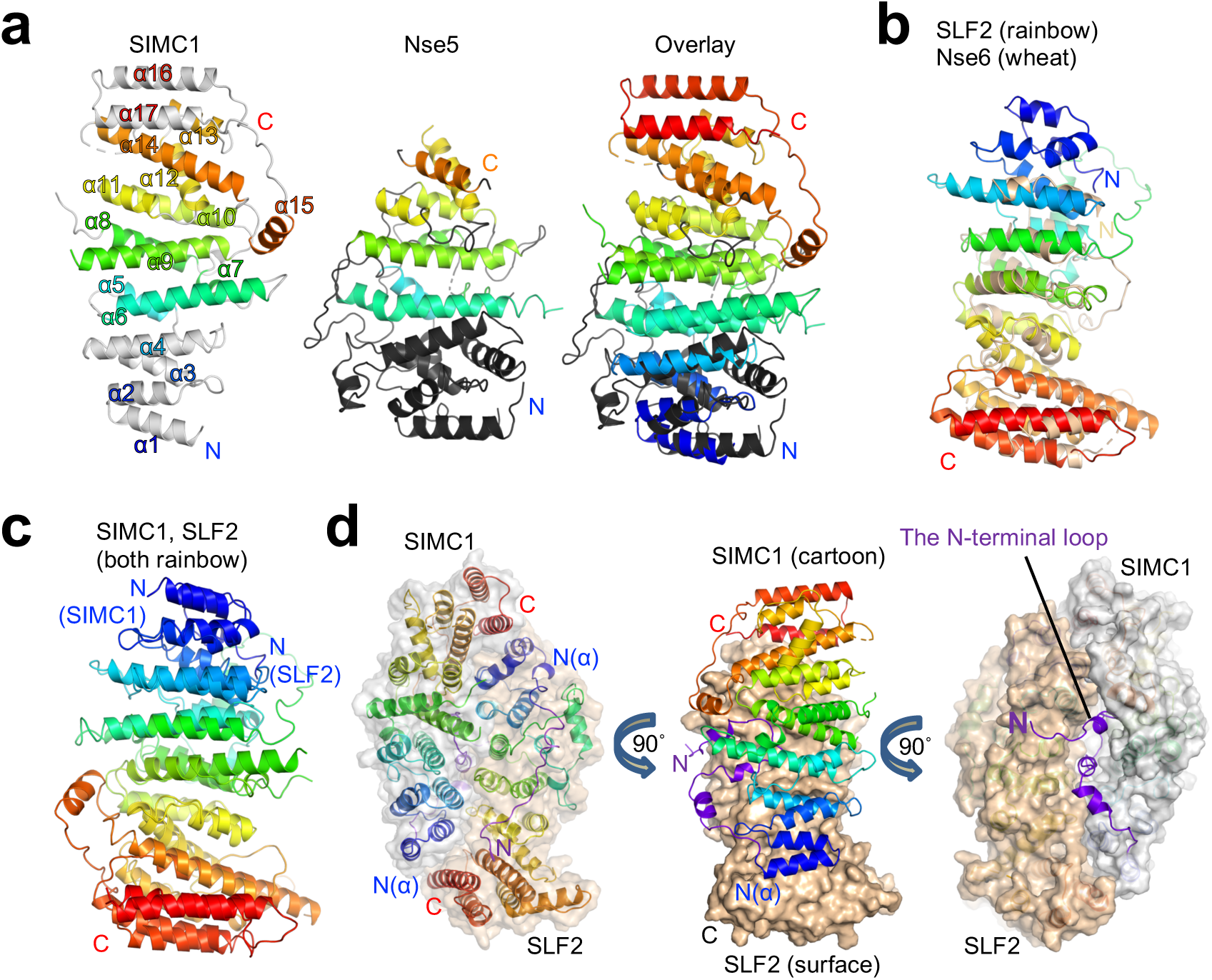
Structural analyses of the SIMC1-SLF2 complex. **(a)** Remote homology mapping on the α-solenoid domain of SIMC1 AlphaFold model (residues 468-858 excluding the 629-653 loop for clarity, left) and ScNse5 structure (PDB ID: 7LTO, middle). The HHpred matches are indicated by the same color. The right panel shows the overlay of SIMC1 and Nse5 wherein SIMC1 is colored by the blue-red full rainbow spectrum and Nse5 is colored by the same manner as in the middle panel. The labels “N” and “C” indicate the N- and C-termini, respectively, of the protein chains. **(b)** Overlay of the α-solenoid domain of SLF2 AlphaFold model (residues 755-1158) and ScNse6 structure (PDB ID:7LTO). **(c)** Overlay of SIMC1 and SLF2 AlphaFold models. **(d)** Cryo-EM structure of the SIMC1-SLF2 complex. Both SIMC1 and SLF2 structures contain a loop-like region at their N-termini, which are colored purple. The label “N(α)” indicates the N-terminus of an α-solenoid fold. On the right panel, SIMC1’s surface is drawn only for its α-solenoid domain to clarify the N-terminal loop of SIMC1.

AlphaFold reveals that the C-terminal Nse6-like region of SLF2 also adopts an α-solenoid-like structure. Although an HHpred search with SLF2 did not identify ScNse6 (**Figure 2 – figure supplement 1**), automated superimposition aligned the SLF2 and ScNse6 structures well, with an r.m.s.d. of 5.7 Å between the structurally aligned 128 Cα atoms (**Figure 5b**). The overlay suggests that SLF2 and Nse6 structures are very similar except for a few α-helices in the N-terminus of SLF2 that are missing in ScNse6. We noted that the SIMC1 and SLF2 α-solenoids are also similar; they can be superimposed well from their N- to C-termini (an r.m.s.d. of 5.6 Å for 247 Cα atoms, **Figure 5c**). However, the similarity between SIMC1 and SLF2 appears to be limited to topology; their sequences do not align, not even remotely, as HHpred fails to detect each other. ScNse5 and ScNse6 structures are too remote to superimpose.

Next, we attempted to predict the structure of the SIMC1-SLF2 complex using ColabFold and AlphaFold-Multimer (47, 48). Both programs yielded the same structure with high confidence (**Figure 5 – figure supplement 2**). To evaluate the accuracy of the predicted structures, we performed cryo-EM single-particle analysis on the SIMC1^284^-SLF2^CTD^ complex (**Figure 5 – figure supplement 3, 4** and **Table 1**). 3D reconstruction yielded an anisotropic and of low-quality map, resolving side-chain densities of only bulky residues in the protein core. The limited orientations and alignability of the particles and the dynamic nature in the end regions of the α-solenoids together likely hampered the reconstruction. Nonetheless, the predicted SIMC1-SLF2 model could be docked into the map unambiguously and refined to ∼3.9 Å resolution. The refined structure was strikingly similar to the predictions, with r.m.s.d.s of only ∼1 Å for Cα atoms (**Figure 5 – figure supplement 5**), validating the accuracy of the predicted structures.

**Table 1.** Cryo-EM data collection/processing and model refinement statistics.

|  |  |  |
| --- | --- | --- |
| Sample | Human SIMC1 (284-872)-SLF2 (635-1173) (EMD-25706, PDB 7T5P) |  |
| Data collection and processing |  |  |
| Condition | 0.4 mg/ml protein | 0.8 mg/ml protein with 0.128 mM DDM |
| Magnification | 73,000 × | 73,000 × |
| Voltage (kV) | 200 | 200 |
| Electron exposure (e<sup>-</sup>/Å<sup>2</sup>) | 66.8 | 66.9 |
| Number of frames per image | 46 | 46 |
| Defocus range (μm) | -0.5 to -1.8 | -0.5 to -1.8 |
| Pixel size (Å) | 0.567 | 0.567 |
| Symmetry imposed | C1 | C1 |
| Initial particle images (no.) | 1,873,621 | 1,065,073 |
| Final particle images (no.) | 31,383 | 8,443 |
| Map resolution (Å) | 3.4 |  |
| FSC threshold | 0.143 |  |
| Refinement |  |  |
| Initial model used | AlphaFold model |  |
| Map resolution (Å<sup>2</sup>) | 3.9 |  |
| FSC threshold | 0.5 |  |
| Map sharpening B factor (Å<sup>2</sup>) | – |  |
| Model composition |  |  |
| Chains | 2 |  |
| Atoms | 13597 (Hydrogens: 6866) |  |
| Protein residues | 822 |  |
| Ligands | 0 |  |
| B-factors (Å<sup>2</sup>) min/max/mean |  |  |
| Protein | 99.26/213.18/135.62 |  |
| Ligand | – |  |
| R.m.s. deviations |  |  |
| Bond lengths (Å<sup>2</sup>) | 0.003 |  |
| Bond angles (°) | 0.591 |  |
| Validation |  |  |
| MolProbity score | 2.00 |  |
| Clashscore | 15.16 |  |
| Rotamer outliers (%) | 0.26 |  |
| Ramachandran plot |  |  |
| Favored (%) | 95.43 |  |
| Allowed (%) | 4.57 |  |
| Disallowed (%) | 0.00 |  |

The cryo-EM structure contains 425-858 aa of SIMC1 and 733-1158 aa of SLF2. SIMC1 and SLF2 α-solenoids face each other in a head-to-tail fashion through the concave surfaces of their slightly curved shapes (**Figure 5d**). Overall, this ellipsoid-shaped structure resembles the yeast Nse5/6 structure (**Figure 5 -figure supplement 6**, (27, 29)). An exception is the most N-terminal region (425-467 aa) of SIMC1 preceding its α-solenoid that adopts an extended conformation. A loop region (431-451 aa) within it is inserted into a cleft between the curved SIMC1 and SLF2 α-solenoids (**Figure 5d**). The loop contributes directly to the formation of the SIMC1-SLF2 interface; without this loop, the α-solenoids’ interface would bury a surface area of only ∼1500 Å^2^, but the complete interface including the loop buries ∼2500 Å^2^. Consistently, our domain mapping experiments showed that the SIMC1 construct (457-872 a.a.) containing the entire α-solenoid but lacking the loop binds SLF2 more weakly than constructs containing the loop (**Figure 2c**). For comparison, the Nse5/6 interface has an open cleft between the subunits (**Figure 5 - figure supplement 6**). However, with the interface burying ∼1800 Å^2^, Nse5 and Nse6 can form a stable complex (20,22,27,29).

The interaction surfaces of both SIMC1 and SLF2 are highly charged (**Figure 5 - figure supplement 7a**). The charges are distributed such that electrostatic interactions strengthen the interface. Furthermore, the interface is enriched with conserved residues (**Figure 5 - figure supplement 7b**), supporting the evolutionary conservation of the architecture from yeast to human. Altogether, these structural features indicate that the SIMC1-SLF2 complex is built as a stable Nse5/6-like heterodimer.

### SIMC1 regulates SMC5/6 localization via its Nse5-like domain

The ellipsoid-like overall shape of the SIMC1-SLF2 complex suggests that the complex likely uses its surface to regulate SMC5/6 function. To explore functional sites, we returned to conservation analysis and found a composite patch consisting of conserved residues of the N-terminus of SIMC1 and the C-terminus of SLF2 (**Figure 6a**). As described above, this end of the complex is structurally preserved between yeast and human complexes. Therefore, this patch may play a role in SMC5/6 regulation.

**Figure 6.**
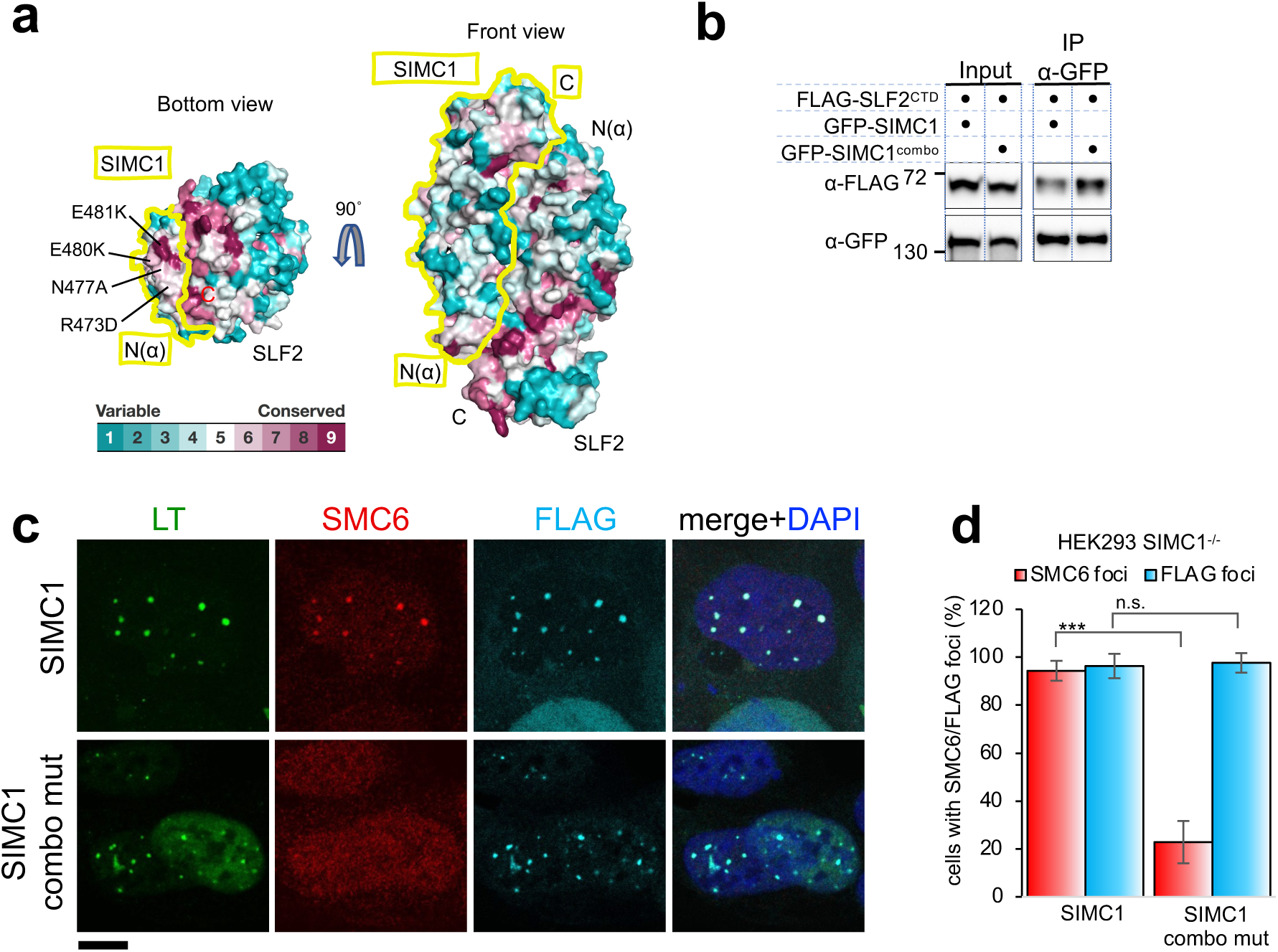
SIMC1 Nse5-like domain regulates SMC5/6 localization at PyVRCs. **(a)** Conservation mapping on the surface of the SIMC1-SLF2 complex. The conservation score obtained from Consurf server are shown by the color graduation as indicated. The boundary of SIMC1’s surface is indicated by yellow lines for clarity. Mutated amino acids (R473D/N477A/E480K/E481K) in the α1 of SIMC1 α-solenoid are labeled. **(b)** Western blot of GFP-trap immunoprecipitation from HEK293 cells transfected with either GFP-SIMC1 or GFP-SIMC1 combo mutant in combination with FLAG-SLF2^CTD^. Full and unedited blots provided in Figure 6-source data 1 **(c)** Representative immunofluorescence images of HEK293 SIMC1^-/-^ cells with integrated vectors expressing FLAG-SIMC1 or FLAG-SIMC1 combo mut, respectively. Cells were fixed 48h after SV40 transfection and stained with SV40 LT (green), SMC6 (red), FLAG (light blue) antibodies. Scale bar 10 μm. **(d)** Relative quantification of the number of cells containing SMC6 and FLAG foci (representative images shown in panel **c**). A minimum of 210 cells with at least four SV40 LT foci were counted for each cell line. Means and error bars (s.d.) were derived from 3 independent SV40 transfections representing biological replicates. (∗) P < 0.05; (∗∗) P < 0.005; (∗∗∗) P < 0.0005; (n.s.) P > 0.05 (two-tailed unpaired t-test). Primary data provided in Figure 6-source data 2.

To test this, we took advantage of our SIMC1 null cells and expressed a epitope-tagged SIMC1 “combo” mutant that contains four mutations (R473D/N477A/E480K/E481K) in the α1 of SIMC1 α-solenoid (**Figure 6a**). These mutation sites are all exposed to solvent and do not make contacts with SLF2. Consistent with this, the SIMC1 combo mutant retained SLF2 binding as demonstrated by co-immunoprecipitation (**Figure 6b**). The SIMC1 combo mutant expressed at a similar level to wild-type SIMC1 and colocalized with LT (**Figure 6b, c**). However, LT localization of SMC5/6 was markedly reduced with the combo mutant compared to wild-type SIMC1 (∼80% reduction) (**Figure 6c, d**). Thus, these data indicate that the Nse5-like domain of SIMC1 is responsible for SMC5/6 recruitment but not for LT localization.

### SLF1 interacts with SLF2 through its split Nse5-like Domain

The sequence of SLF1, the putative human orthologue of yeast Nse5, lacks detectable Nse5-like regions but contains an ankyrin repeat domain (ARD) (33, 39). We turned to the AlphaFold model of SLF1, which reveals a flexible architecture containing BRCT domains at the N-terminus and the ARD and an α-solenoid-like helical domain in the C-terminus (SLF1^CTD^; 410-1058 aa, **Figure 7a**). Notably, the ARD (726-935 aa) is an insert of the α-solenoid, which strikingly resembles SIMC1’s Nse5-like domain, with an r.m.s.d. of 5.9 Å between the structurally aligned 300 Cα atoms (**Figure 7b**). We then ran an HHpred search with a C-terminal SLF1 sequence lacking the ARD (SLF1^CTD_ΔARD^), which identified Nse5s and SNI1 (**Figure 7 – figure supplement 1a**). Thus, SLF1 does contain an Nse5-like domain split by an ARD in its sequence.

**Figure 7:**
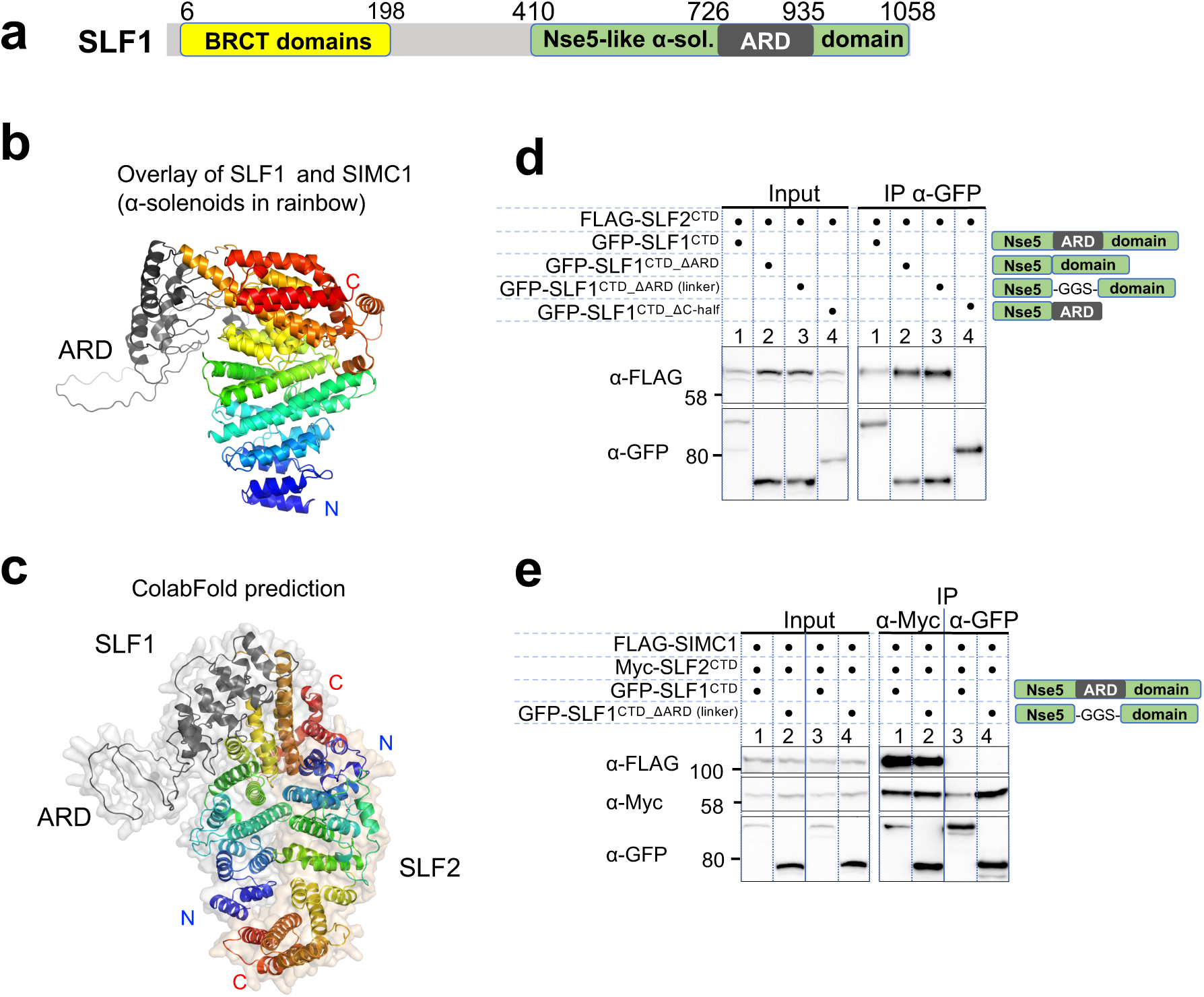
SLF1 interacts with SLF2 through its split Nse5-like domain. **(a)** Schematic of SLF1 domain boundaries on the AlphaFold model. **(b)** Superposition of the α-solenoid domains of SLF1 and SIMC1. **(c)** The ColabFold-predicted structural model of the SLF1^CTD^-SLF2 complex. **(d)** Western blot of GFP-Trap immunoprecipitation from HEK293 cells transiently transfected with plasmids expressing the respective protein combination. Schematic on the right represents domains in SLF1 truncated variants. Full and unedited blots provided in Figure 7-source data 1 **(e)** Western blot of Myc-Trap or GFP-Trap immunoprecipitation from HEK293 cells transfected with FLAG-SIMC1, Myc-SLF2^CTD^ and GFP-SLF1^CTD^ or GFP-SLF1^CTD_ΔARD^ ^(linker)^, respectively. Domains of SLF1 constructs are shown in the scheme on the right side. Full and unedited blots provided in Figure 7-source data 2.

We then asked ColabFold for an SLF1^CTD^-SLF2 structure, which predicted with high confidence a model similar to the SIMC1-SLF2 structure (**Figure 7c, Figure 7 – figure supplement 2a**). Because the ARD does not make contact with SLF2, we repeated the prediction with SLF1^CTD_ΔARD^, which produced the same structure (**Figure 7 – figure supplement 2b**). Predictions using a short SLF1 lacking the C-terminal half of the split Nse5-like domain (SLF1^CTD_ΔC-half^; 410-935 aa) and another SLF1 containing only the N-terminal half of the split Nse5-like domain (SLF1^CTD_N-half^; 410-725 aa) failed to assemble a reliable complex (**Figure 7 – figure supplement 2c, d**). These results suggest that the complete Nse5-like domain would be necessary and sufficient for the association of SLF1 with SLF2.

We then experimentally verified that expressed SLF1^CTD^ binds to the SLF2^CTD^ (**Figure 7d**). Consistent with the *in silico* results, SLF1^CTD_ΔARD^ retained interaction with SLF2^CTD^, but SLF1^CTD_ΔC-half^ did not do so (**Figure 7d**). Next, we asked if SIMC1 and SLF1 form exclusive complexes with SLF2. SIMC1, SLF1 and SLF2 were co-expressed in cells and subjected to immunoprecipitation of either SLF1 or SLF2. Whilst both SIMC1 and SLF1 are found in SLF2 immunoprecipitates, only SLF2 was detected following immunoprecipitation of SLF1 (**Figure 7e**). Therefore, either SLF1 or SIMC1 forms an Nse5/6-like complex with SLF2, which agrees with the structural data showing that SLF1 and SIMC1 bind to the same surface on SLF2. The mutually exclusive binding of SIMC1 and SLF1 to SLF2 is further supported by recent proteome-wide mass spectrometry analyses (**Figure 7 – figure supplement 3**, (17, 49)).

We wondered if the structural and functional similarities between SIMC1 and SLF1 are encoded in the primary sequence. To test this, we ran an HHpred search with the SLF1 sequence against the human proteome, which identified SIMC1 (**Figure 7 – figure supplement 1b**). To validate the HHpred result, we manually generated a sequence alignment between SIMC1 and SLF1 based on the structural alignment described above (**Figure 7 – figure supplement 4**) and compared it with the HHpred alignment (**Figure 7 – figure supplement 5**). While the registrations are slightly off from the structure-based alignments, the HHpred alignment matches most of the structurally corresponding helices, covering from α1 to α14, which is the first helix of the C-terminal half of the split Nse5 domain in the SLF1 sequence. Altogether, our results establish structural and functional (SLF2 binding) similarities between SLF1 and SIMC1. We conclude that the two proteins are distant paralogues.

## Discussion

Our study identifies SIMC1 as an elusive human orthologue of yeast Nse5, establishes that SLF1 is also an Nse5 orthologue, and reveals evolutionary specialization of these key regulators of SMC5/6 function. SIMC1 and SLF1 form exclusive complexes with the Nse6 ortholog SLF2. Like Nse5/6, SIMC1-SLF2 and SLF1/2 function as SMC5/6 recruiters, and notably, they play distinct roles (**Figure 8**). We showed that SIMC1-SLF2 localizes SMC5/6 to PyVRCs, whereas SLF1/2 is known to recruit SMC5/6 to DNA lesions (33). The different readers of post translational modifications (PTMs) in each protein likely mediate such distinct targeting.

**Figure 8:**
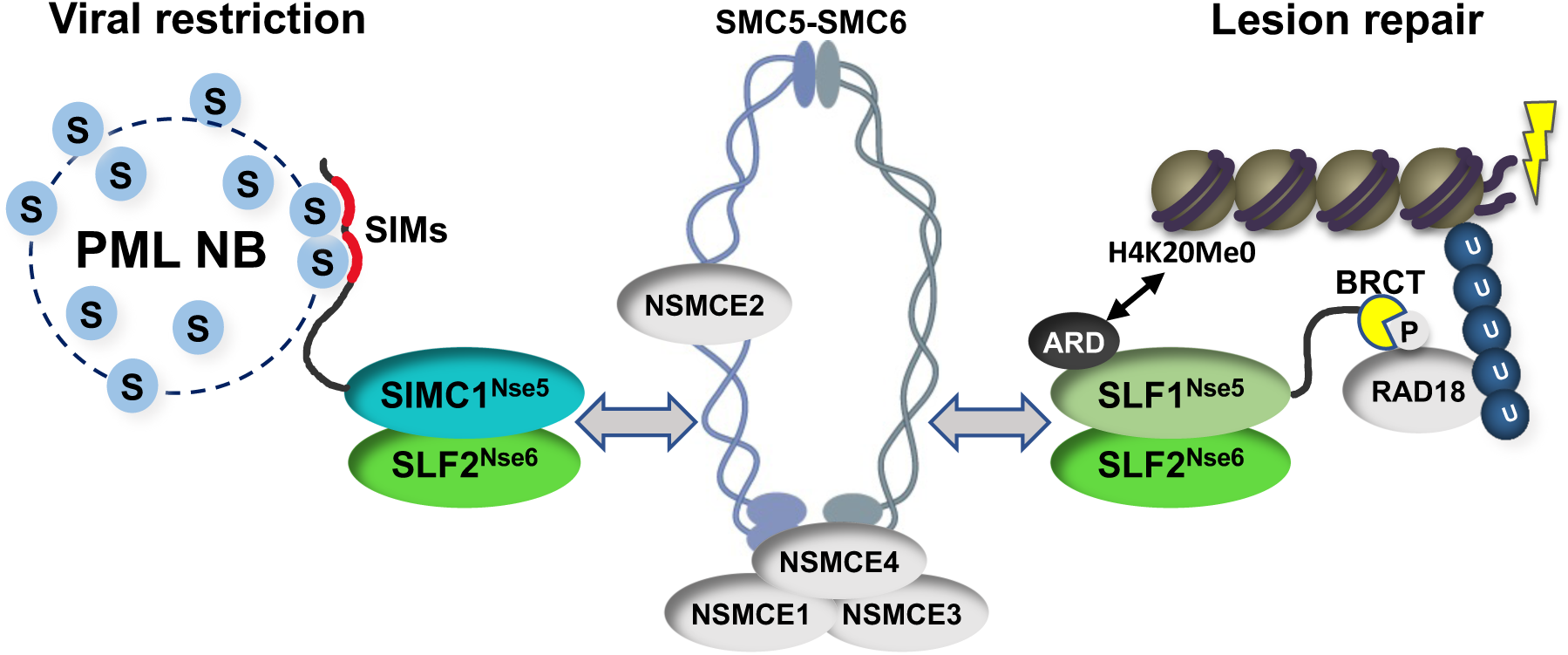
Schematic of SIMC1-SLF2 and SLF1/2 complex function. Human orthologues of yeast Nse5, SIMC1 and SLF1, form exclusive complexes via their C-terminal Nse5-like domains with the human Nse6 ortholog SLF2. The SIMC1-SLF2 complex localizes SMC5/6 to SV40 replication centers, whereas SLF1/2 targets it to DNA lesions, likely via recognition of different post translational modifications. S stands for SUMO; U for ubiquitin.

Proteins use tandem SIMs to recognize SUMOylated proteins (35) and enter SUMO-rich PML NBs via multivalent SIM-SUMO interactions (50). SIMC1 also relies on its SIMs to localize at PML NBs, thereby mediating the enrichment of SMC5/6 at PyVRCs. Once at viral replication centers, SIMC1, like yeast Nse5, may promote the SUMO ligase activity of SMC5/6 (23,24,29,51), which could in turn reinforce SIMC1-mediated SMC5/6 recruitment (**Figure 8**). Here, the interaction of SIMC1 with the poly-SUMO2 ligase ZNF451 could help generate SUMO2 chains (42, 43) that are in turn recognized by the SIM motifs of SIMC1.

SLF1 uses its N-terminal BRCT domains to bind to the DNA repair protein RAD18 at DNA lesions (33). This mechanism parallels the recruitment of Smc5/6 to chromatin in yeast (23,24,30). Brc1/Rtt107 uses its BRCT domains to bind Rad18 and gamma-H2A at DNA lesions, while it also binds Nse6 within the Nse5/6 complex. Thus, by inclusion of BRCT domains, SLF1 appears to have combined Nse5 and Brc1/Rtt107 functions (**Figure 8**). SLF1 also uses its ARD to bind the unmethylated histone H4 tail (H4K20Me0), which is only found on nascent chromatin (52).

Complete understanding of the SMC5/6 recruitment mechanisms by SIMC1-SLF2 and SLF1/2 requires further work, but our data indicate that SIMC1’s Nse5-like domain is involved. In addition, SLF2 has been reported to interact directly with SMC6 (39). Thus, it is likely that SIMC1 or SLF1 and SLF2 together bridge the recruitment PTMs (e.g., SUMO, phospho-RAD18 and H4K20Me0) and the SMC5/6 core (**Figure 8**). Supporting this model, both Nse5 and Nse6 are essential for the recruitment of Smc5/6 to chromatin in yeast, and both contact Smc5/6 in crosslinking-MS studies (20,23,24,26,27,29,31). Nse5/6 also regulates Smc5/6 function by inhibiting its ATPase activity (26, 27). Thus, the SIMC1-SLF2 and SLF1/2 complexes may act similarly to direct SMC5/6 activity.

Our discovery of SIMC1-dependent localization of SMC5/6 to PML NBs suggests a broad role of SMC5/6 in the antiviral response. Many viral genomes including HBV, HSV-1, HPV, SV40 and the pathogenic human polyomaviruses e.g., JCV are deposited next to, or seed the formation of, PML NBs (53–58). There are pro- and antiviral properties ascribed to PML NBs, but it is generally accepted that these bodies and some of the proteins they contain pose a barrier to productive infection by several viruses (53-56,58). In the case of HBV the restrictive roles of SMC5/6 and PML NBs are intertwined, as SMC5/6 transcriptionally silences HBV cccDNA only when localized at PML NBs (53). This is likely a more pervasive functional pairing given the localization of many viruses to PML NBs and the increasing number of these shown to be antagonized by SMC5/6 (11,13,15,17). For example, SIMC1 expression is induced ∼7 fold upon expression of the early HPV E7 protein in HaCaT cells (59), and recent work suggests that SMC5/6 restricts high-risk HPV (15). Therefore, SIMC1-SLF2 may recruit SMC5/6 to antagonize the replication of HBV, HPV and other viruses.

The list of viruses SMC5/6 antagonizes is growing and has recently acquired HIV-1 (17). In this case, SLF2 was shown to recruit SMC5/6 to, and restrict the expression of, non-integrated HIV-1 genomes, but these effects occurred in the absence SLF1. Given our discovery reported here, it is tempting to speculate that SIMC1 supports this SMC5/6 function. Such a case would strengthen the notion that SIMC1 is more specialized toward antiviral function. Further characterization of SIMC1 will be a promising strategy to establish the role of SMC5/6 in fighting pathogenic viruses and uncover the underlying molecular mechanisms.

## Materials and Methods

### Construction of recombinant plasmids

Plasmid DNA was constructed either by standard restriction enzyme digestion and T4 ligase ligation-based cloning or using Takara In-Fusion HD Cloning Plus Kit. To make SIMC1 SIM mutations (FIDL to AADA and VIDL to AADA, corresponding to SIMC1 amino acids 26 to 30, and 45 to 49), a gblock sequence containing all mutation was purchased (IDT). Combo mutations in SIMC1 Nse5-like domain were introduced in primers and two overlapping fragments were cloned into a plasmid using In-Fusion kit. All plasmids created in this study have been verified by sequencing service provided by Eton Bioscience (San Diego). Plasmids and oligonucleotides used in this study are provided (**Supplementary file 3**). Additional details of plasmid construction are available upon request.

### Cell culture, transfection, stable line generation

Human embryonic kidney cell lines HEK293 and HEK293T, human osteosarcoma cell line U2OS, and Phoenix ampho cells were obtained from American Type Culture Collection (ATCC). Flp-In™-293 cell line was purchased from ThermoFisher (R75007). Cells were cultured in Dulbecco’s modified Eagle’s medium (DMEM) supplemented with 10% fetal bovine serum (FBS).

Transient plasmid transfections were generally carried out using TransIT-LT1 (Mirus) transfection reagent or Polyethylenemine (Polysciences Inc.) at a 3:1 transfection reagent to DNA ratio. To generate stable cell lines with ectopic expression from plasmids, lentivirus or retrovirus was produced in HEK293T or Phoenix ampho cells, respectively, and used to infect target cell lines, followed by drug selection. Flp-In™-293 cell lines were generated by co-transfecting 0.4 µg of the relevant FRT construct (**Supplementary file 3**) and 3.6 µg of pOG44 (ThermoFisher) in 10 µl of Lipofectamine 2000 (ThermoFisher) diluted in 0.5 mL of OptiMEM (ThermoFisher), following the manufacturer’s instructions. Hygromycin B was added 2 h after transfection and selection continued until colonies started to appear.

A complete list of stable mammalian cell lines generated and reagents used in this study is provided (**Supplementary file 3**).

### Cell line generation by CRISPR/Cas9

SIMC1 knockout clones were generated via CRISPR/Cas9 gene targeting by transient transfection of a hSpCas9 encoding plasmid (Addgene) and a pcDNA-H1 plasmid encoding specific sgRNA: guide1 (5’-GGGTCTGAACGACATAACGC-3’), guide 2 (5’-CGCAGGAAAAGGACTCGCCC-3’), both in exon 5. To introduce stop codons by HR-mediated repair, a donor template with the STOP cassette sequence (GTCGGATCCTTTAAACCTTAATTAAGCTGTTGTAG) was used. The presence of the STOP cassette in clones from single cell were confirmed by sequencing, depletion of SIMC1 was verified by RT-qPCR and western blot. For RT-qPCR, total RNA was precipitated using RNeasy Plus Mini kit (Quiagen), cDNA synthetized by SuperScript III First-Strand Synthesis System for RT-PCR (Invitrogen) and SensiFAST SYBR No-ROX kit (Meridian Bioscience) was used for qPCR.

### SILAC SMC5 BioID labeling and affinity purification of biotinylated proteins

For stable isotope labeling by amino acids in cell culture (SILAC), 8 × 10^6^ cells were seeded in a 15-cm plate with 45 mL of DMEM for SILAC (ThermoFisher) supplemented with foetal calf serum (10% v/v), 2 mM L-glutamine, 100 U/mL penicillin-streptomycin and the relevant amino acids: arginine-0/lysine-0 (light, 84 and 146 mg/L, Sigma-Aldrich, A6969 and L8662), arginine-6/lysine-8 (heavy, Cambridge Isotope Laboratories, CNLM-539-H-1 and CNLM-291-H-1). After 48 h, biotin (stock: 50 mM in DMSO) was added to the medium to a final concentration of 50 µM for BirA*-SMC5 or 0.0125 µM for BirA*, and cells were cultured in the presence of biotin for 24 h. Cells were detached by trypsinization, then counted and diluted to ∼ 1.25 × 10^8^ in 50 mL of PBS. Equal volumes of light and heavy-labelled cells were mixed as follows: 25 mL of BirA*-light with 25 mL of BirA*-SMC5-heavy and 25 mL of BirA*-heavy with 25 mL BirA*-SMC5-light. Cells were centrifuged at 300 *g* for 5 min, then incubated on ice for 15 min in 30 mL of 10 mM HEPES pH 7.6, 25 mM NaCl, 1.5 mM MgCl_2_, 0.34 M sucrose, 10% v/v glycerol, 1× cOmplete protease inhibitors, 1 mM DTT (buffer A), supplemented with 0.1% v/v Triton X-100. The released nuclei were harvested by centrifugation at 1500 *g* for 5 min, washed once in 30 mL of buffer A supplemented with 0.1% v/v Triton X-100 and twice more in 30 mL of buffer A without Triton X-100. Purified nuclei were suspended in 2 mL of 1% v/v SDS, 10 mM EDTA pH 8, 20 mM HEPES pH 7.6, 1× cOmplete protease inhibitors (SDS buffer) and boiled for 10 min. Nuclear lysate was sonicated using a Branson 450 Sonifer equipped with a cone micro tip in five pulses of 15-20 s at minimum power settings to reduce sample viscosity. Nuclear lysate was diluted with 8 mL of 25 mM HEPES Ph 7.6, 150 mM NaCl, 1 mM EDTA pH 8, 1× cOmplete protease inhibitors, and 1 mM DTT (dilution/wash buffer), and incubated overnight at 4° C with 400 µl of MyOne Streptavidin T1 Dynabeads that was pre-washed in dilution buffer. The beads were washed three times with 10 mL of dilution/wash buffer, and eluted twice with 20 µl of 1× NuPAGE LDS Sample Buffer supplemented with 100 mM of DTT and 1 mM of biotin for 5 min at 95° C.

### Sample preparation and mass spectrometry of SMC5 BioID with SILAC

The immunoprecipitated samples were run on the 10% SDS-PAGE for 10 min at 180V to allow the proteins to move into the resolving gel. The gel was then fixed for 15 min in 7% acetic acid and 40% methanol solution and stained for 15 min with 0.25% Coomassie Blue G-250, 7% acetic acid, and 45% ethanol. After rinsing with deionized water to remove excess dye, In-gel digestion was performed in principle as described previously (60) and detailed as follows. Each gel lane was cut separately using the new sharp scalpel, then minced, and transferred to an Eppendorf tube. Gel pieces were destained with 50% ethanol and 25 mM NH_4_HCO_3_ for 15 min to remove Coomassie dye. After removing supernatant, the gel pieces were dehydrated with 100% acetonitrile for 10 min on the rotator. Acetonitrile was discarded, and the samples were dried to completion using a vacuum evaporator (Eppendorf). The dried samples were then rehydrated and disulphide bonds in the proteins were reduced using 10 mM DTT in 50 mM NH_4_HCO_3_ pH 8.0 (reduction buffer) for 1 h at 56°C. The buffer was removed, and cysteine residues of proteins were subsequently alkylated with 50 mM iodoacetamide and 50 mM NH_4_HCO_3_ pH 8.0 for 45 min at room temperature in dark. Samples were dehydrated again with 100% acetonitrile, then dried by vacuum evaporation. The fully dried gel slices were incubated with 1 µg trypsin per tube in 50 mM ammonium bicarbonate buffer at 37°C overnight on a ThermoMixer (Eppendorf). Digested peptides were extracted twice with 150 µl of 30% acetonitrile, then with 150 µl of 100% acetonitrile to the gel pieces for 15 min at 25°C while being agitated at 1400 rpm in a ThermoMixer (Eppendorf). Extracted peptides combined. The reductive dimethylation step was performed as described previously (61). The labelled samples (each sample pair including a replicate were switched with the SILAC (Lys8/Arg10) labels) were mixed, and purified and desalted with C18 Stage-Tips (M3 company) as described (62). The eluted peptides were loaded on the silica column of 75 µm inner diameter (New Objective) packed to 25 cm length with 1.9 µm, C18 Reprosil beads (Dr. Maisch Phases) using an Easy-nLC1000 Liquid Chromatography system (Thermo Scientific).

Peptides were separated on the C18 column using an Easy-nLC1000 Liquid Chromatography system (Thermo Scientific) with the following 2 h reversed-phase chromatography gradient: 0-4 min, 2-5% solvent B; 4-67 min, 5-22% solvent B; 67-88 min, 22-40% solvent B; 88-92 min, 40-95% solvent B; 92-97 min, 95% solvent B; 97-101 min, 95-2% solvent B; and 101-105 min, 2% solvent B (solvent B: 80% acetonitrile containing 0.1% formic acid) and directly sprayed into a Q-Exactive Plus mass spectrometer (Thermo Scientific) for the data acquisition. The mass spectrometer was operated in the positive ion scan mode with a full scan resolution of 70,000; AGC target 3×10^6^ max. IT = 20 ms; Scan range 300 - 1650 m/z with a top10 MS/MS DDA method. Normalized collision energy was set to 25 and MS/MS scan mode operated at a resolution of 17,000; AGC target 1×10^5^ and max IT of 120 ms.

### Nano-Trap immunoprecipitation

GFP and Myc-labeled proteins were immunoprecipitated using the Nano-Trap magnetic agarose (Chromotek) following the manufacturer’s instructions. Briefly, about 10 x10^6^ cells were harvested by trypsinization 48-72 h after plasmids transfection, lysed by incubation in 200 μl of dilution buffer (10 mM Tris pH 7.5, 150 mM NaCl, 0.5 mM EDTA) supplemented with 0.5% Nonidet NP40, 1 mM phenylmethylsulfonyl fluoride (PMSF), 1x Halt Protease Inhibitor Cocktail (Thermo Fisher Scientific) for 30 min on ice with extensive pipetting every 10 min. Lysates were cleared by centrifugation at 17,000 *g* 12 min and input aliquot was incubated with 6 mM MgCl_2_ and Benzonase (1:75, EMD Millipore). The remaining lysate was diluted with 300 μl of dilution buffer + supplements and incubated with 25 μl of GFP- or Myc-Trap magnetic agarose beads for 2 h at 4°C, followed by three washes in dilution buffer and elution into 2x NuPAGE LDS Sample Buffer (ThermoFisher). Samples were analyzed in SDS-PAGE and western blot.

### SIMC1 BioID labeling and affinity purification of biotinylated proteins

BioID labeling was carried out essentially as described (63). Briefly, cells were cultured in media containing 50 μM biotin (Sigma) for 24 h. Denaturing cell lysis and streptavidin pulldown was performed as described previously (38, 63). Native lysis purification was carried out as detailed below. Cell pellet was suspended in native lysis buffer (20 mM Tris, pH 7.5, 150 mM NaCl, 1 mM EDTA, 0.5% NP-40, 1 mM DTT, 6 mM MgCl_2_, 1 mM PMSF) supplemented with Halt Protease Inhibitor Cocktail (ThermoFisher). The cell lysate was incubated on ice for 30 min with 50 units of Benzonase (EMD Millipore), then centrifuged at 4°C at 16,000 *g* for 10 min. Supernatant, diluted with 3 volumes of 10 mM Tris, pH 7.5, 150 mM NaCl (native dilution buffer), was incubated for 2 h with Dynabeads MyOne Streptavidin C1 beads (ThermoFisher, 20 μl of beads was used in small-scale analysis; 75 μl in large scale preparation for mass spectrometry). The beads were then washed three times with 10 mM Tris, pH 7.5, 500 mM NaCl (native wash buffer). Following washes, beads were either eluted with 35 μl of 2x NuPAGE LDS Sample Loading Buffer (ThermoFisher) with 100 mM DTT at 100°C for 5 min for Western blotting, or stored in 100 mL of 8M urea, 100 mM Tris, pH 8.5, for mass spectrometry analysis.

### Protein identification by mass spectrometry

Protein samples were reduced and alkylated by sequential incubation with 5 mM Tris (2-carboxyethyl) phosphine for 20 min at room temperature and 10 mM iodoacetamide reagent in the dark at room temperature for additional 20 min. Proteins were digested sequentially at 37°C with lys-C for 4 h followed by trypsin for 12 hours. After quenching the digest by the addition of formic acid to 5% (v/v), peptides were desalted using Pierce C18 tips (Thermo Fisher Scientific), dried by vacuum centrifugation, and resuspended in 5% formic acid. Peptides were fractionated online using reversed phase chromatography on in-house packed C18 columns as described before (64). The 140 min gradient of increasing acetonitrile was delivered using a Dionex Ultimate 3000 UHPLC system (Thermo Fisher Scientific). Peptides were electrosprayed into the mass spectrometer by the application of a distal *2.2 kV* spray voltage. MS/MS data were acquired using an Orbitrap Fusion Lumos mass spectrometer operating in Data-Dependent Acquisition (DDA) mode consisting of a full MS1 scan to identify peptide precursors that were subsequently targeted by MS2 scans (Resolution = 15,000) using high energy collision dissociation for the remainder of the 3 second cycle time. Data analysis was performed using the Integrated Proteomics bioinformatic pipeline 2 (Integrated Proteomics Applications, San Diego, CA). Database searching was performed using the ProLuCID algorithm against the EMBL Human reference proteome (UP000005640 9606). Peptides identifications were filtered using a 1% FDR as estimated using a decoy database. Proteins were considered present in a sample if they had 2 or more unique peptides mapping to them. Relative comparisons between samples to identify candidate SIMC1 interacting proteins was done using raw peptide spectral counts.

### Immunofluorescence and microscopy

Immunofluorescence was performed essentially as described (63). Briefly, U2OS cells grown on coverslips were fixed in 4% formaldehyde in PBS for 10 min, then permeabilized with PBS containing 0.2% Triton X-100 for 5 min at room temperature. The cells were incubated with antibodies diluted in a blocking solution of 5% Normal Goat Serum (BioLegend) in PBS. HEK293 cells were grown on coverslips coated in 0.1% gelatin (type A, MP bio), fixed in 4% formaldehyde in PBS for 15 min, permeabilized in Triton X-100 buffer (0.5% Triton X-100, 20 mM Hepes, 50 mM NaCl, 3 mM MgCl_2_, 300 mM sucrose) for 2 min. Antibodies were diluted in PBS containing 0.2% cold water fish gelatin (Sigma) and 0.5% BSA.

To visualize DNA, fixed cells were stained with 0.1 μg/mL 4’,6-diamidino-2-phenylindole dihydrochloride (DAPI, Sigma). Coverslips were mounted in ProLong Gold Antifade (Invitrogen) to glass slides. Antibodies used for immunofluorescence are listed (**Supplementary file 3**).

Confocal images were acquired with LSM 780 or 880 confocal laser scanning microscope (Zeiss) equipped with 63x/1.4 oil immersion high NA objective lens, using standard settings in Zen software. Images were processed using ImageJ software (NIH).

### Western blotting

Whole cell lysate was prepared by lysis cells in RIPA buffer (150 mM NaCl, 1% Triton X-100, 0.5% sodium deoxycholate, 0.1% SDS, 50 mM Tris, pH 8), agitated for 40 min in 4°C and centrifuged at 16,000 *g* for 20 min. The whole cell lysate or pulldown sample was combined with NuPAGE LDS Sample Loading Buffer and 100 mM DTT, separated by SDS-PAGE and transferred to nitrocellulose. Immunoblotting was performed as described (65). Antibodies used for immunoblotting are listed (**Supplementary file 3**). Protein detection was carried out using ECL chemiluminescent substrates (genDEPOT, ThermoFisher) on a ChemiDoc XRS+ Molecular Imager (BioRad) using ImageLab software (BioRad).

### Flow cytometry

HEK293 cells were treated with 100 ng/mL nocodazole in 0.1% DMSO for 10 h to induce G2/M arrest. Cells were fixed with ice-cold 70% EtOH, washed in PBS containing 1% BSA and stained with 50 μg/mL propidium iodide in the presence of 250 μg/mL RNAse A for 30 min in 37°C and 4°C overnight. Flow cytometry analysis was performed on a NovoCyte 3000 (ACEA Biosciences) using NovoExpress software.

### Alkaline comet assay

Endogenous level of DNA damage in HEK293 cells was evaluated using the CometAssay kit (Trevigen) according the manufacturer’s instructions. Control cells were treated with 1 mM MMS (Sigma) for 1 h to induce DNA damage. Cells were harvested by trypsinization and 500 cells were mixed with LMAgarose and spread on the slide. Slides were dried 20 min in 4°C, incubated 40 min in kit lysis solution in 4°C and 20 min in unwinding solution (300 mM NaOH, 1 mM EDTA, pH 13) in room temperature. Electrophoresis was performed in the unwinding solution in 4°C, 300 mA, 30 min. Slides were washed twice in water, once in 70% EtOH, dried 15 min in 37°C and stained with SYBR Gold (Thermo) for 30 min. Images were taken using Zeiss Axio Imager MI with 20x objective.

### Recombinant protein expression and purification

Recombinant human SIMC1 and SLF2 proteins were expressed in baculovirus-infected Spodoptera frugiperda (Sf9) insect cells. The cDNA fragment encoding SIMC1^284^ (284-872 aa) was placed after the Tev-protease cleavable GST tag in a modified pACEBac vector. The cDNA fragment encoding SLF2^CTD^ (a.a. 635-1173) was cloned after the Tev-cleavable Histidine tag in a modified pACEBac. Baculoviruses were generated from these vectors using the published protocol (66). Sf9 cells were co-infected with the baculoviruses harboring GST-SIMC1^284^ or His-SLF2^CTD^ and harvested after 48-50 h. The cells were suspended in the lysis buffer (20 mM Hepes pH 7.5, 150 mM NaCl, 0.5 mM TCEP) supplemented with protease inhibitor cocktail (Pierce) and lysed using C3 pressure homogenizer (Avestin). The lysate was clarified by centrifugation at 40,000 *g*. Imidazole (pH 7) was added to the supernatant, which was then poured onto a Ni affinity column (Thermo Fisher Scientific). The Ni resin was washed with five column volume of the lysis buffer. Proteins were eluted with 300 mM Imidazole. The Ni eluate was immediately poured onto a glutathione sepharose column (GoldBio). After wash, an aliquot of Tev protease was added to the resin, which was then left at 4°C overnight. Proteins were then eluted in the lysis buffer, concentrated, and injected a Superdex 200 size exclusion column (GE Healthcare). The protein contents of the fractions were analyzed by SDS-PAGE, which was stained with Coomassie blue.

### ColabFold and AlphaFold-Multimer Structural modeling

ColabFold and AlphaFold-Multimer were run on Google Colab platform using default settings.

### Cryo-EM data collection and processing

Cryo-EM grids were prepared in cold room. A 3 µl drop of the SIMC1^284^-SLF2^CTD^ complex (0.4 mg/ml protein in 20 mM HEPES pH 7.5, 150 mM NaCl, 0.5 mM TCEP or 0.8 mg/ml protein in 20 mM HEPES pH 7.5, 150 mM NaCl, 0.5 mM TCEP, 0.128 mM DDM) was applied to a plasma-cleaned UltraAuFoil R1.2/1.3 300-mesh grids (Quantifoil). Immediately after removing excess liquid by a filter paper (Whatman No. 1), the grid was rapidly dropped into liquid ethane using a manual plunger. Grids were stored in liquid nitrogen tank until use.

Data were collected in two sessions, which were done in a same manner as follows. Grids were observed on Talos Arctica 200 KeV transmission electron microscope (Thermo Fisher Scientific) equipped with a K2 Summit direct detector (Gatan). The microscope was aligned as reported previously (67). Images were recorded in an automated manner using the Leginon software (68). Data were collected as movie files with a frame rate of 200 ms in the counting mode. The acquisition parameters are described in Table 1. Particle motion correction was performed using the MotionCor2 software (69) in the Appion data processing pipeline (70). The frame-aligned images were imported into Relion 4.1b (71) and CTF estimation was performed using gCTF (72). The first session on the condition of 0.4 mg/ml protein concentration collected 3332 micrographs. The grids contained many well-frozen areas, which enabled an efficient data collection. A total of 1,873,621 particles were picked from the first data set using Laplacian-of-Gaussian filter. Particle images were extracted at 2.268 Å per pixel with a box size of 100 pixels and subjected to 2D classification. 2D classes showing secondary structures were selected and subjected to *ab initio* reconstruction in cryoSPARC (73), which yielded a model that appears to be consistent with 2D class average images. The *ab initio* model was 3D classified in Relion with K=3 and T=4. The particles belonging to the highest resolution model (620,042 particles, 41.0 % of the total input particles) were selected and 3D auto-refined in Relion. The auto-refined particles were re-centered and re-extracted at 1.134 Å per pixel with a box size of 200 pixels, then were subjected to 2D classification. 203,464 particles were selected and subjected to 3D auto-refinement. The refined particles were combined with the particles from the second data set as described below.

Processing of the first data set made it clear that the orientation of the particles was limited. To alleviate this issue, DDM (0.75 × critical micelle concentration) was added to the sample and the concentration of the protein was increased. The images of this sample appeared to show more different views of the particles; however, the grid contained too many empty holes in areas with thin ice, which made it difficult to obtain a large number of good particles. As such, the second session collected 1553 micrographs. A total of 1,065,073 particles were picked using Laplacian-of-Gaussian filter, extracted at 2.268 Å per pixel with a box size of 100 pixels, and then subjected to 2D classification. 128,371 particles belonging to well-resolved 2D class average images were selected and 3D auto-refined. The refined particles were re-centered and re-extracted at 1.134 Å per pixel with a box size of 200 pixels, and then subjected to 3D auto-refinement. The refined particles were 3D classified with no alignment with K=2 and T=8. 30932 particles belonging the better-resolved class (24.2 % of the total input particles) were selected and 3D auto-refined. The particles were then combined with the refined particles from the first data set, yielding a total of 234,396 particles. The combined particles were subjected to 3D auto-refinement, followed by 3D classification without alignment (K=2, T=12). 39,826 particles belonging to the better-resolved map (17.0 % of the total input) were 3D auto-refined. The refined particles were subjected to CTF refinement, but no improvement was observed. Local resolution variation was calculated in Relion. Three-dimensional FSC analyses were performed on the remote 3DFSC processing server (https://3dfsc.salk.edu/, (74)). The final map was sharpened by DeepEMhancer (75) on the COSMIC2 server (https://cosmic-cryoem.org/, (76)).

### Model building, refinement and structural analyses

For model building, a SIMC1-SLF2 model predicted by AlphaFold-Multimer was docked into the sharpened cryo-EM map using the Dock in map module of PHENIX (77). The model was trimmed and adjusted into the map using COOT (78) and refined using the Real-space refinement module of PHENIX. The map-to-model FSC was obtained from PHENIX. The refined model was validated using MolProbity (79). The buried surface areas of the SIMC1-SLF2 (PDB ID: 7T5P) and Nse5/6 (PDB ID: 7LTO) complexes were obtained from ‘Protein interfaces, surfaces and assemblies’ service PISA at the European Bioinformatics Institute (http://www.ebi.ac.uk/pdbe/prot_int/pistart.html, (80)). Electrostatic potential was calculated using APBS (81). Conservation scores were obtained from Consurf server (https://consurf.tau.ac.il, (82). Structure figures were generated using Pymol (https://pymol.org), Chimera (83) and ChimeraX (84).

## Data Availability

The SMC5 and SIMC1 BioID datasets have been deposited to the PRIDE database (85) as follows: Protein interaction AP-MS data: PRIDE PXD033923. Cryo-EM density map and atomic coordinates of the SIMC1-SLF2 complex have been deposited to the Electron Microscopy Data Bank and wwPDB, respectively, under accession codes EMD-25706 and PDB 7T5P.

## Material Availability

All materials produced during this work are available upon written request, in keeping with the requirements of the journal, funding agencies, and The Scripps Research Institute.

## Acknowledgements

We thank Dr. James DeCaprio and Dr. Xiaohua Wu for the kind gift of SV40 plasmids. We thank Chinatsu Otomo for the generation of baculoviruses. We thank Charly Chahwan (Synthex, Inc). for sharing his early insights into Nse5 orthologues. This work was supported by NIH grants R35 GM136273 to M.N.B., GM089788 to J.A.W., and GM092740 to T.O. Computational resource for Cryo-EM data processing was supported by NIH S10OD021634. Funding for the H.D.U. laboratory was provided by the Deutsche Forschungsgemeinschaft (DFG; German Research Foundation; project number 393547839 – SFB 1361, sub-project 07). Support of the IMB Proteomics Core Facility and use of IMB’s Q-Exactive Plus mass spectrometer is also gratefully acknowledged.

## Author information

Contributions

Conceptualization, M.N.B., T.O., E.L.D. Design of Methodology, M.O., M.N., M.N.B., S.M, T.O., J.A.W., Y.J-A., N.Z. Writing – Original Draft, M.N.B., T.O.

Writing – Review & Editing, M.O., M.N., T.O., M.N.B., E.L.D. Formal analysis/Investigation, M.O., M.N., M.N.B., S.M., T.O., J.A.W., Y.J-A., N.Z.,

H.D.U. Validation, M.O., M.N. Resources, M.O., M.N., S.M. Visualization, M.O., M.N., T.O., M.N.B. Funding acquisition, M.N.B., T.O., J.A.W., H.D.U. Data curation, J.A.W., Y.J-A., T.O. Supervision, M.N.B., T.O., J.A.W., H.D.U. Project administration, M.N.B.

Corresponding authors

Correspondence to Michael N. Boddy or Takanori Otomo

## Ethics declarations

Competing interests

The authors declare no competing interests.

## Figure supplements

**Figure 2 - figure supplement 1.**
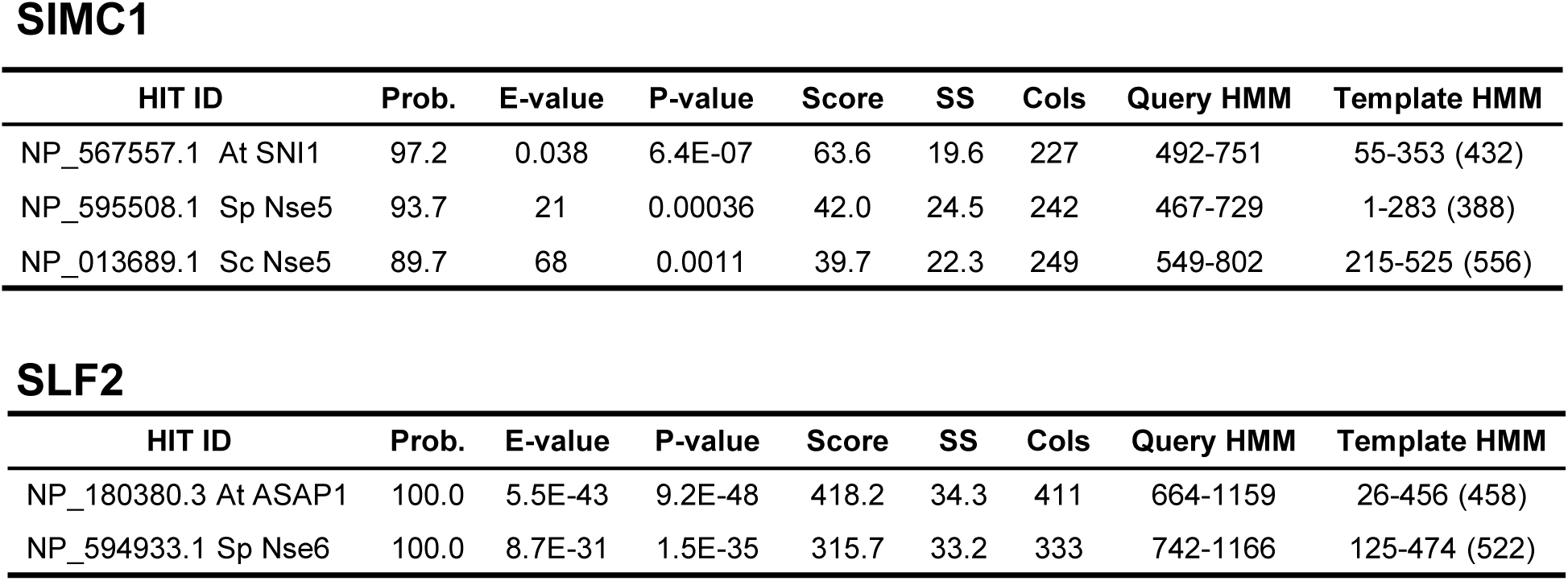
HHPred homology searches for full-length SIMC1 (Q8NDZ2) and SLF2 (Q8IX21). Query proteomes: *Arabidopsis thaliana*, *Saccharomyces cerevisiae*, *Schizosaccharomyces pombe*.

**Figure 2 – figure supplement 2.**
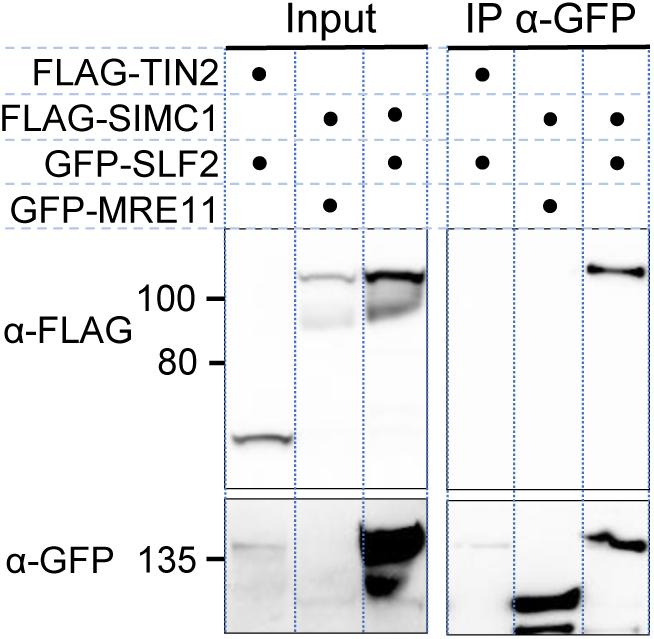
Full-length SIMC1 and SLF2 co-immunoprecipitate. Western blot of GFP-Trap immunoprecipitation from HEK293 cells transfected with plasmids expressing the indicated combinations of proteins. Full and unedited blots provided in Figure 2-figure supplement 2-source data 1.

**Figure 2 – figure supplement 3.**
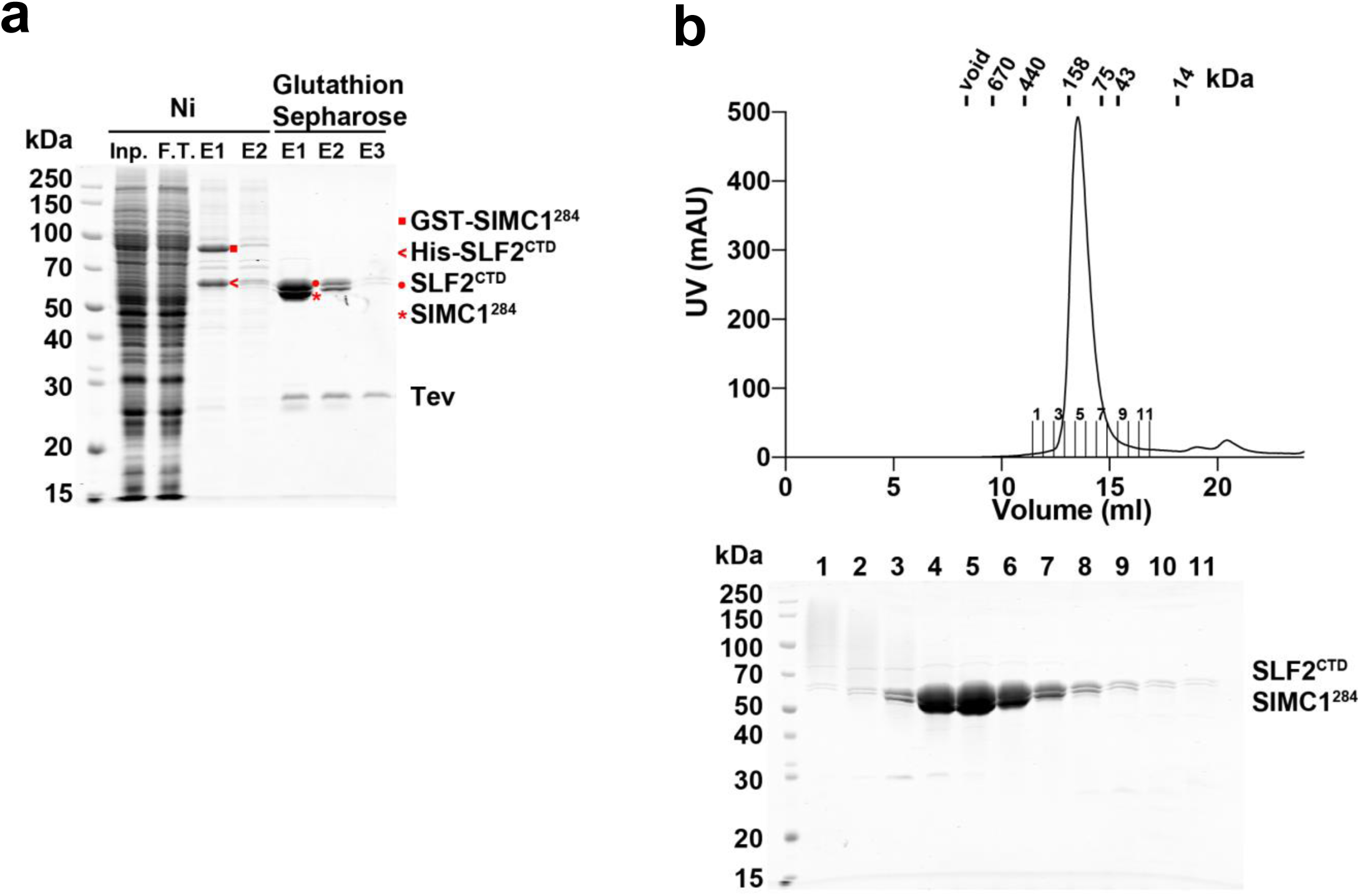
Purification of the SIMC1^284^-SLF2^CTD^ complex from insect cells. **(a)** SDS-PAGE analysis of Ni- and glutathione-affinity purifications. The Ni elution contained GST-SIMC1^284^ and His-SLF2^CTD^. The Ni eluate was loaded onto glutathione sepharose and proteins were eluted by Tev protease digestion. The elution fractions contained untagged SIMC1^284^ and SLF2^CTD^. **(b)** Superdex 200 size exclusion chromatography of the glutathione elution. SIMC1^284^ and SLF2^CTD^ co-eluted in a single peak at the position corresponding to a globular domain of ∼120kDa, which matches with the mass of the complex. Full and unedited gels provided in Figure 2-figure supplement 3- source data 1.

**Figure 4 – figure supplement 1.**
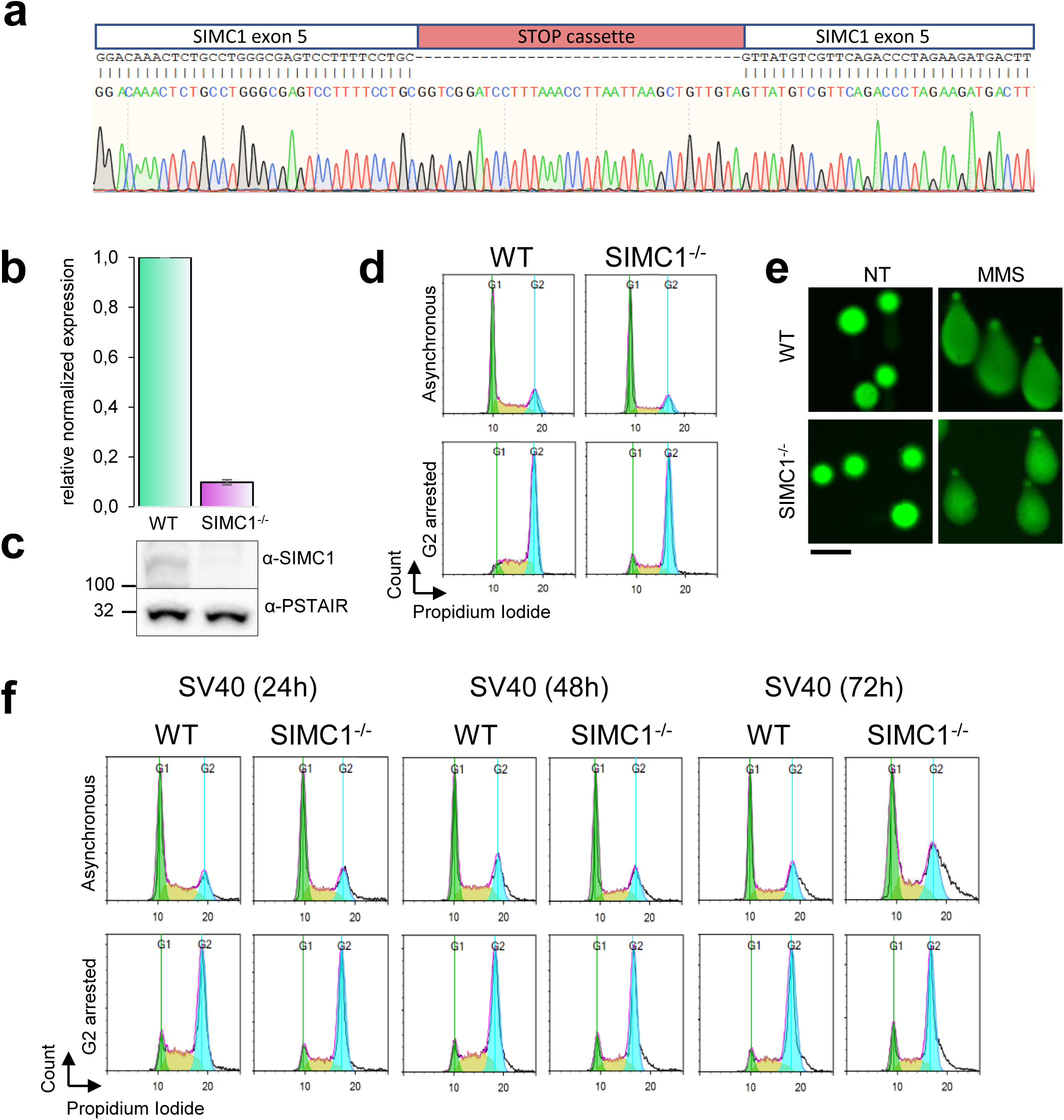
Validation of HEK293 SIMC1-/- clone. **(a)** Sanger sequencing of exon 5 to verify the insertion of STOP cassette (red). **(b, c)** SIMC1 expression was measured by quantitative PCR and normalized to the expression of beta-actin (mean ± s.d. from n=4 independent experiments) and also analyzed on western blot using PSTAIR as a loading control. Primary data for graph in panel **b** and full, unedited blots for panel **c** provided in Figure 4-figure supplement 1-source data 1, 2. **(d)** Flow cytometry analysis of cell cycle profile of asynchronous or G2/M nocodazole arrested cells. **(e)** Comet assay assessing the spontaneous DNA damage. As a control, cells were treated with 1 mM MMS for 1 h. Scale bar 100 μm. **(f)** Flow cytometry analysis of cell cycle profile of asynchronous or nocodazole-treated cells arrested in G2/M in cells transfected with SV40. Cells were treated with nocodazole at the indicated time points after SV40 transfection.

**Figure 5 – figure supplement 1.**
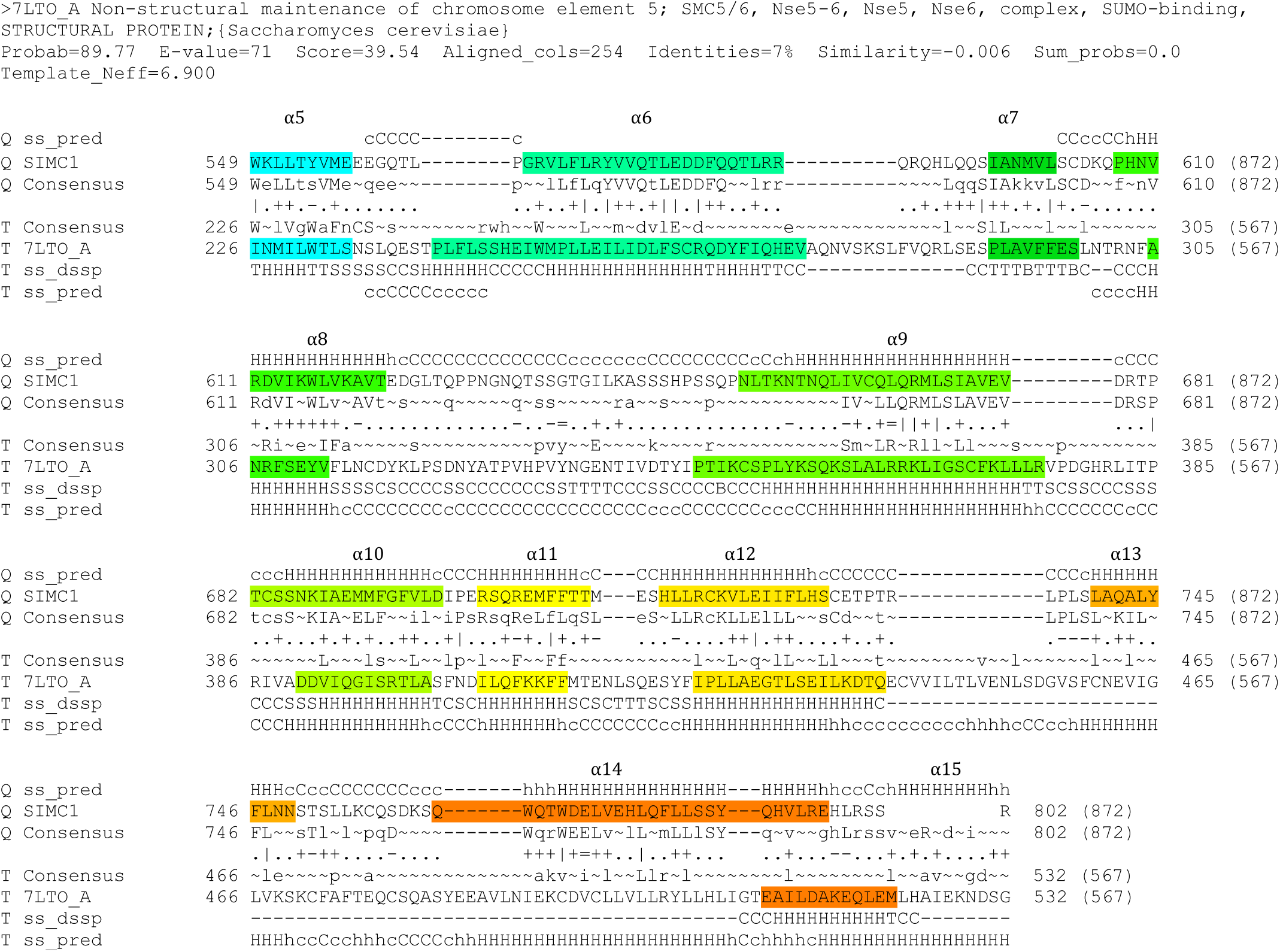
HHpred-aligned SIMC1 and ScNse5 (7LTO_A) sequences. The α-helices determined by experimental structures are colored. The colors match with the rainbow used for cartoon structures shown in Fig. 5.

**Figure 5 – figure supplement 2.**
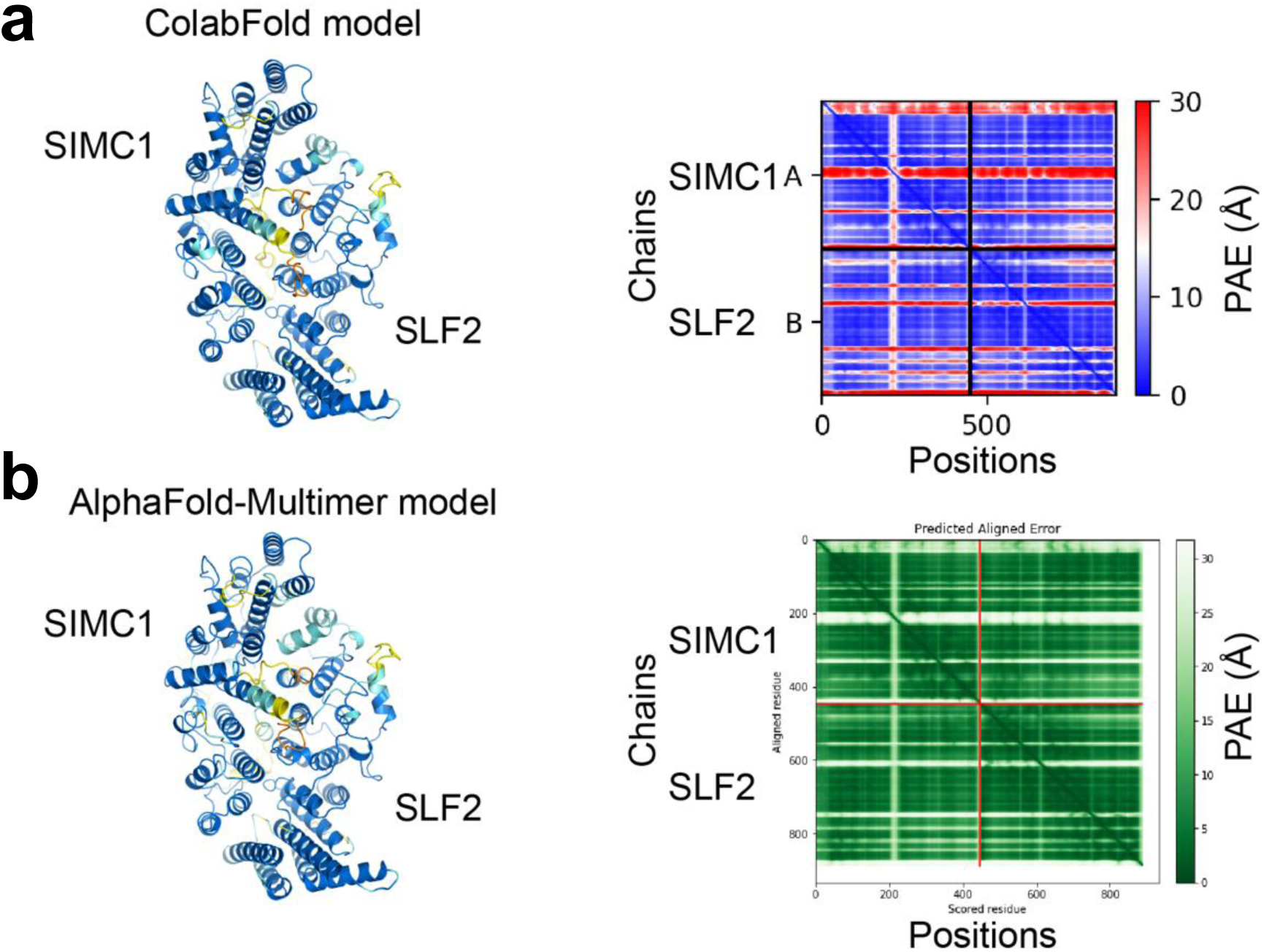
Predictions of the SIMC1-SLF2 structure by ColabFold **(a)** and AlphaFold-Multimer **(b).** SIMC1 (425-872 aa) and SLF2 (733- 1173 aa) were used. The predictions are shown on the left and colored by pLDDT. Orange, yellow, sky blue, and blue indicate very low (pLDDT < 50), low (50 < pLDDT < 70), confident (70 < pLDDT < 90), and high (pLDDT > 90) regions, respectively. PAE plots are shown on the right. Both plots show low PAE values between subunits, indicating high confidence of the predictions.

**Figure 5 – figure supplement 3.**
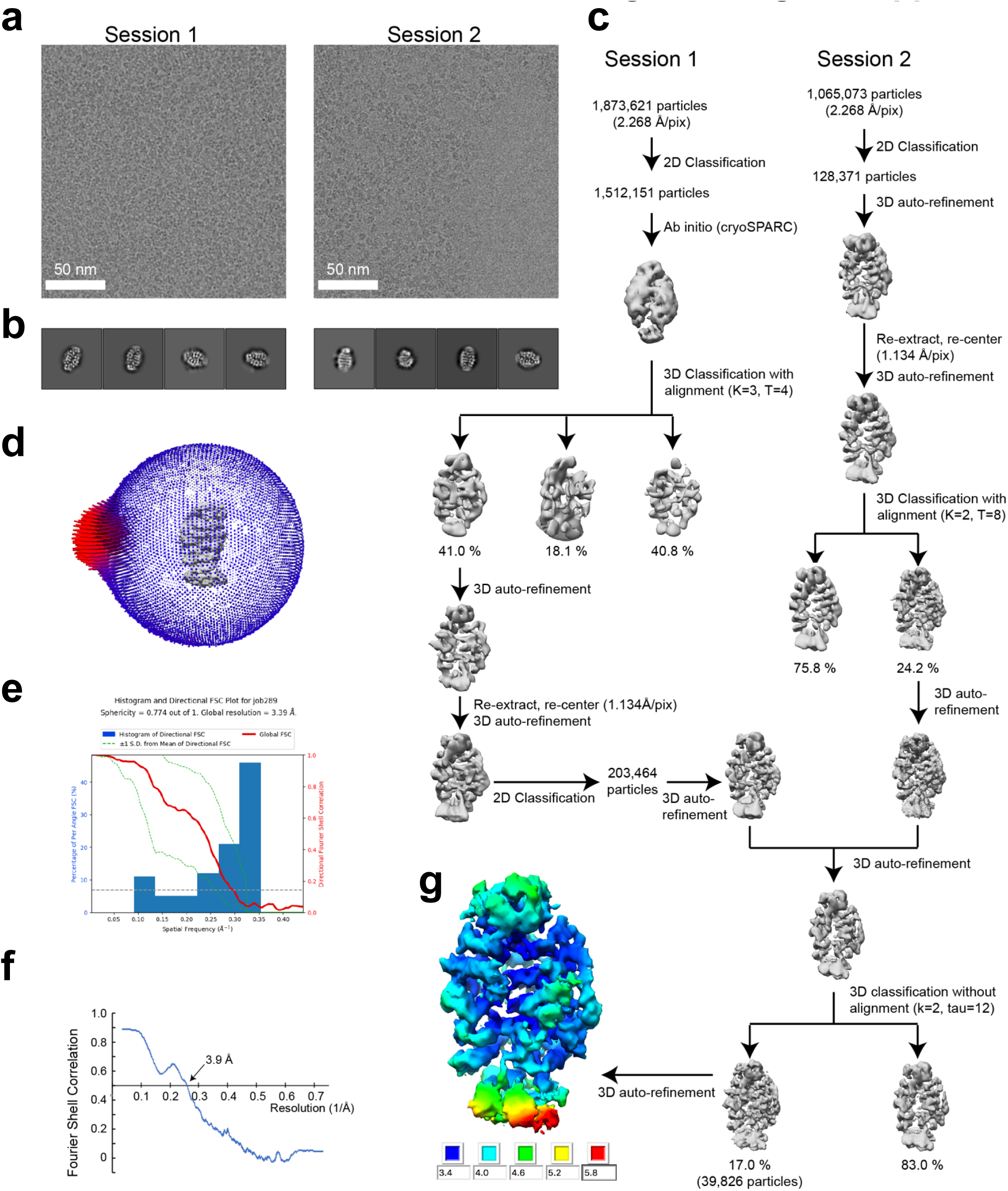
Single-particle cryo-EM data processing of the SIMC1-SLF2 complex. **(a)** Representative cryo-electron micrographs from Session1 and 2. **(b)** Representative 2D class average images of Session 1 and 2. **(c)** Flow chart of 3D reconstruction. **(d)** Euler angle distribution of the final particle images. **(e)** 3D FSC plot. **(f)** Map-to-model FSC plot. **(g)** Local resolution estimation shown on the final map (unsharpened).

**Figure 5 – figure supplement 4.**
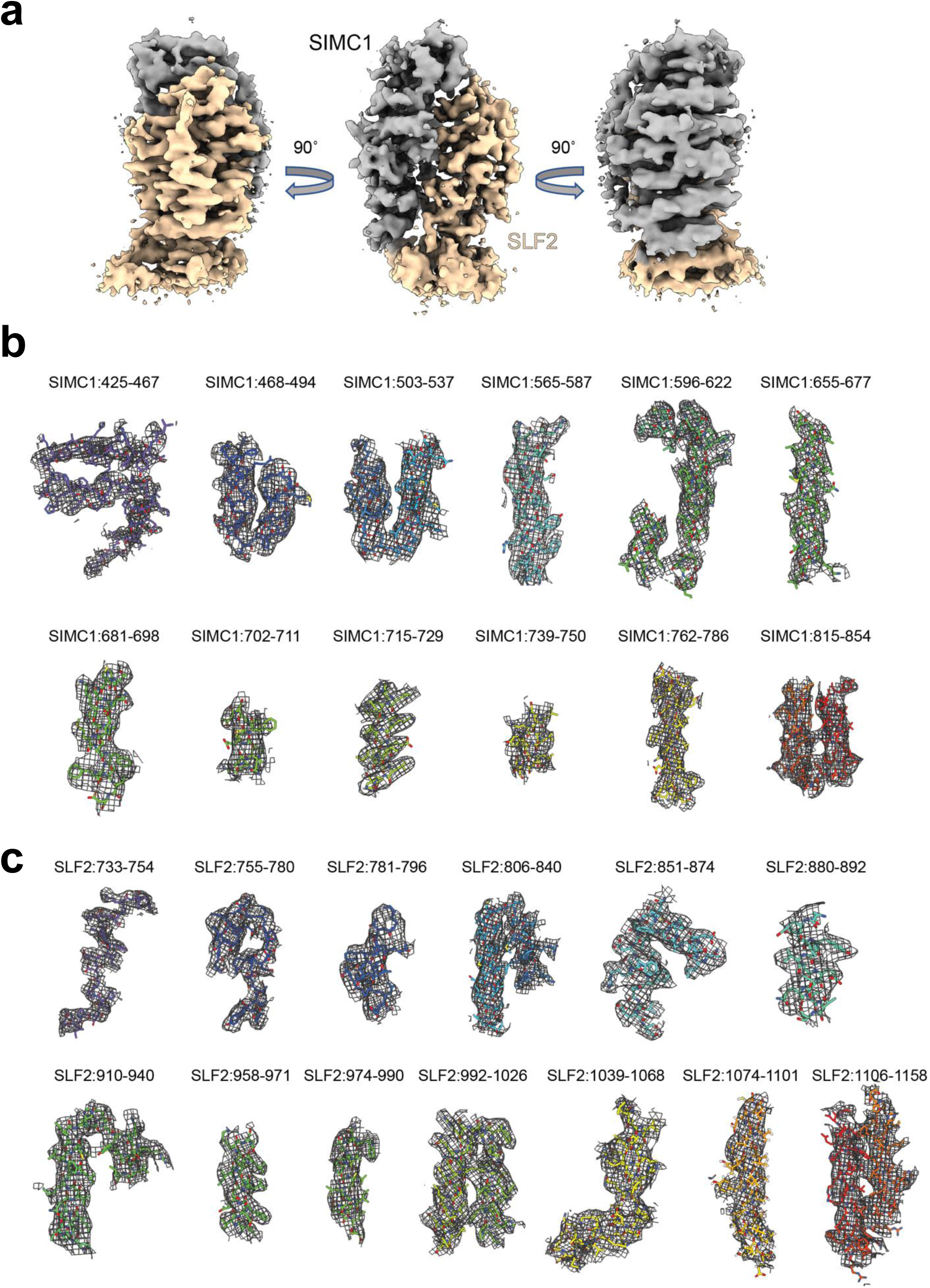
Cryo-EM density map of the SIMC1-SLF2 complex. **(a)** The sharpened map. **(b)** SIMC1 fragments. **(c)** SLF2 fragments. The residue regions are indicated.

**Figure 5 – figure supplement 5.**
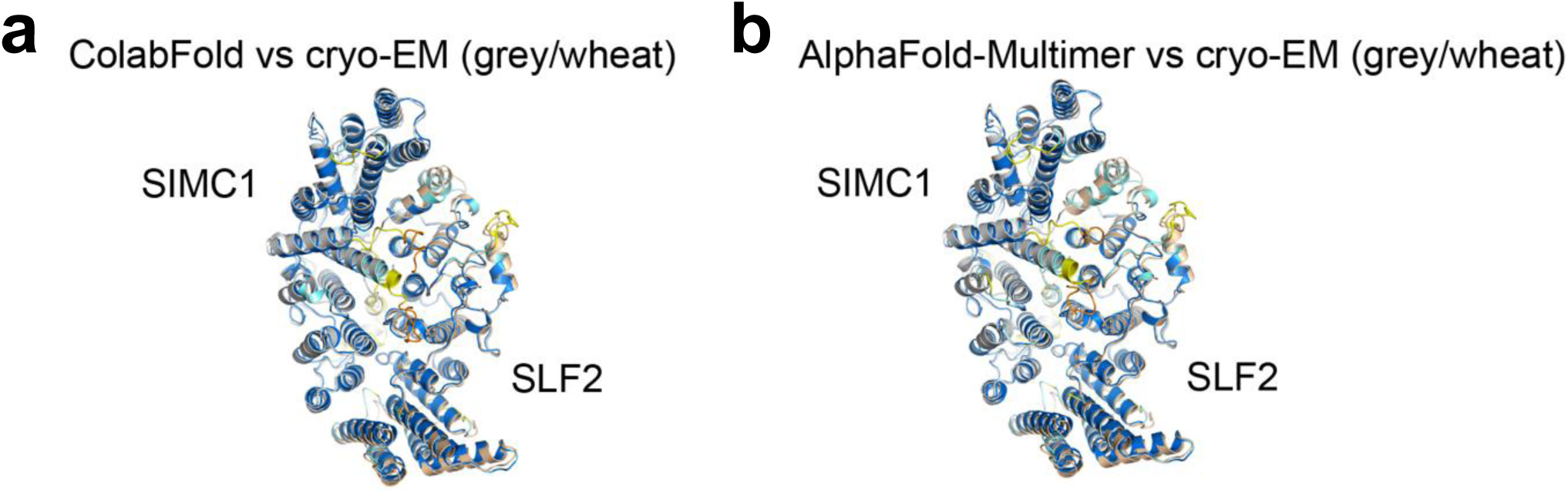
Comparison of the predicted structures and the refined cryo-EM structure. **(a, b)** Structures predicted by ColabFold (a) and AlphaFold Multimer (b) are superimposed on the cryo-EM structure. Prediction structures are colored by pLDDT. SIMC1 and SLF2 of the cryo-EM structure are colored grey and wheat, respectively.

**Figure 5 – figure supplement 6.**
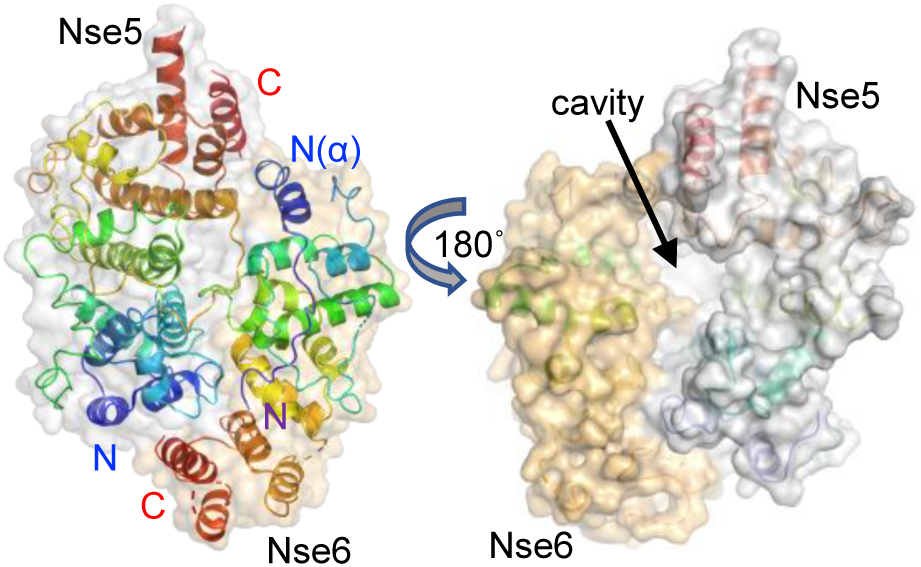
Structure of the ScNse5/6 complex (PDB ID: 7LTO). For comparison with the SIMC-SLF2 structure, ScNse5/6 is shown in a similar orientation as in Fig. 5d.

**Figure 5 – figure supplement 7.**
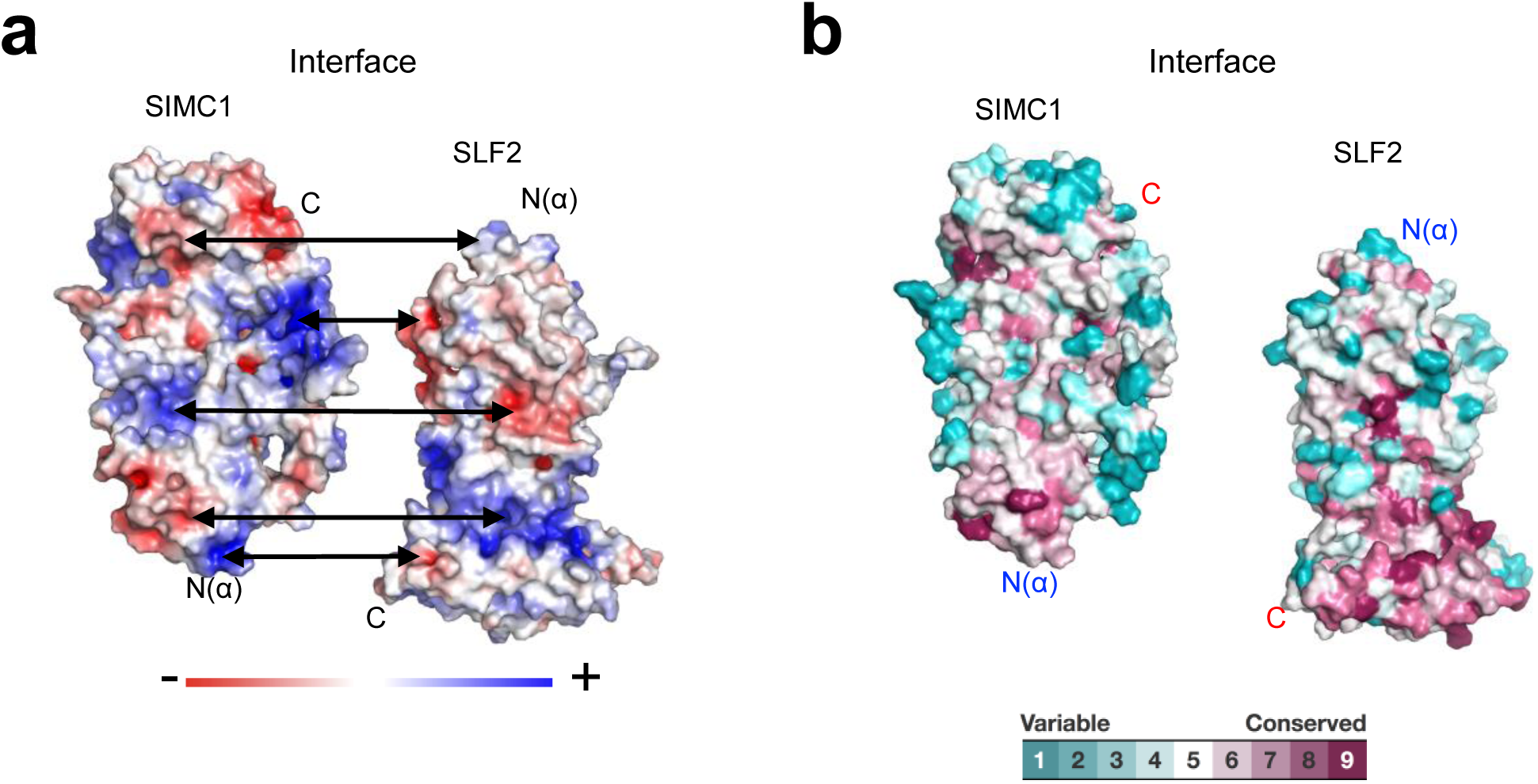
Properties of the SIMC1-SLF2 interface. **(a)** Electrostatics potential of the SIMC1 and SLF2 surfaces. SIMC1 and SLF2 are oriented to show their interfaces. Double arrows point at intramolecular charge pairs. **(b)** Conservation mapping on the subunit interface. The conservation scores obtained from the Consurf server are shown by the color graduation as indicated.

**Figure 7 – figure supplement 1.**
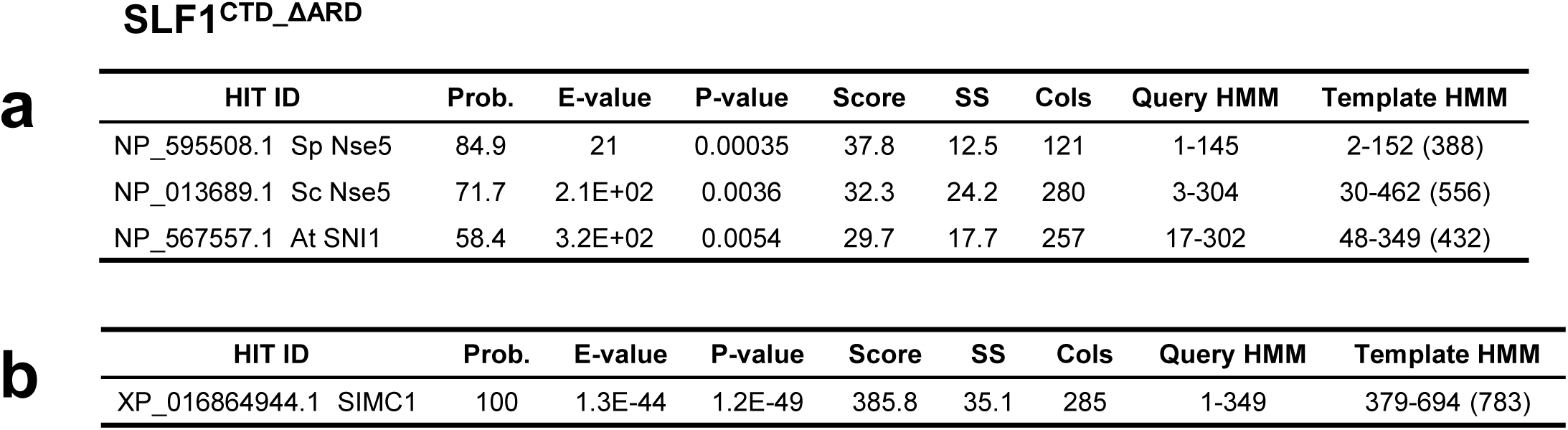
Results of an HHpred search with SLF1^CTD_ΔARD^ (410-725 aa + 935-1058 aa) against yeast and plant sequences (a) and human sequences (b).

**Figure 7 – figure supplement 2.**
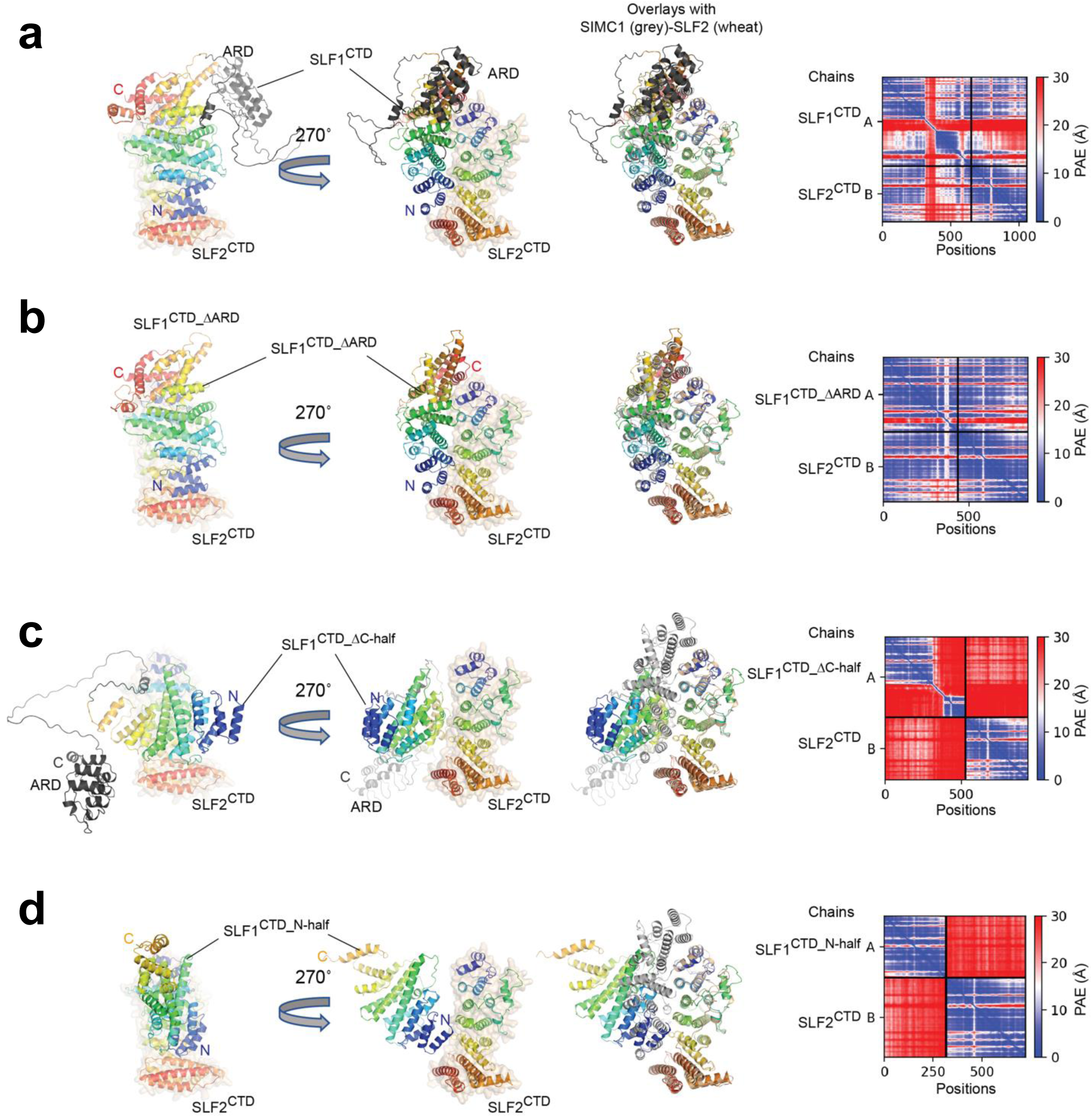
ColabFold structure prediction with SLF1 variants and SLF2^CTD^ (755-1159 aa). **(a)** SLF1^CTD^ (410-1058 aa). **(b)** SLF1^CTD_ΔARD^ (410-725 aa + 935-1058 aa). **(c)** SLF1^CTD_ΔC-half^ (410-935 aa). **(d)** SLF1^CTD_N-half^ (410-725 aa). The best-ranked structure of five predictions is shown for each prediction. Overlays with the SIMC1-SLF2 structure are generated by superimposing SLF2. SIMC1 and SLF2 of the SIMC1-SLF2 structure are colored by grey and wheat, respectively. SLF1 and SLF2 of the predictions are colored in rainbow. The right panels show the PAE plots. The inter-protein PAE values in (a) and (b) are low, indicating high confidence of the predictions, while those in (c) and (d) are high, indicating low confidence of the prediction.

**Figure 7 – figure supplement 3.**
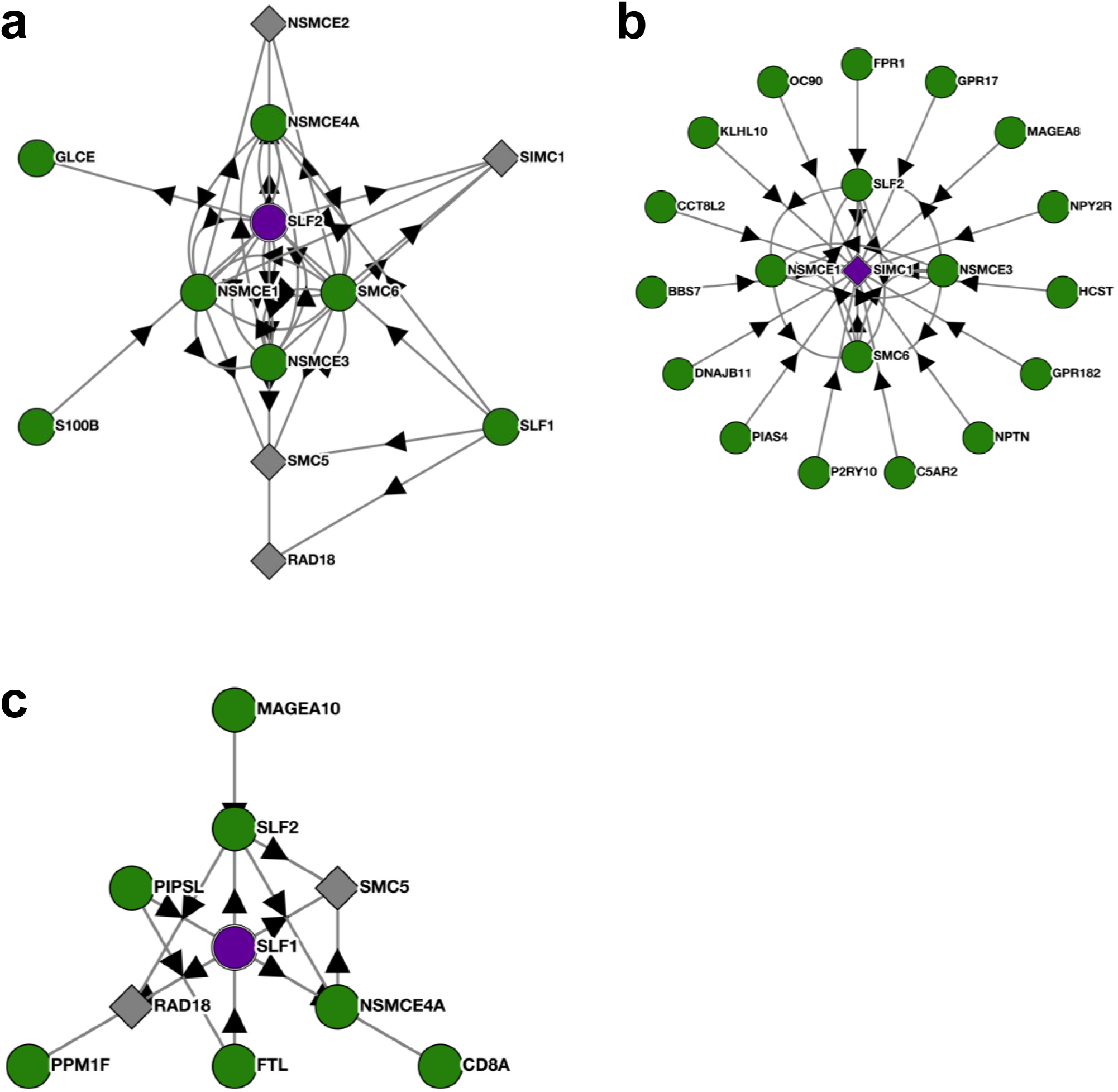
BioPlex interactome nodes for SLF2, SIMC1 and SLF1. **(a-c)** Unedited SLF2, SIMC1 and SLF1 interactome nodes from the Harvard BioPlex Interactome https://bioplex.hms.harvard.edu. The “bait” protein is colored purple at the center of each node, and interacting proteins are shown around the bait with connecting arrows.

**Figure 7 – figure supplement 4.**
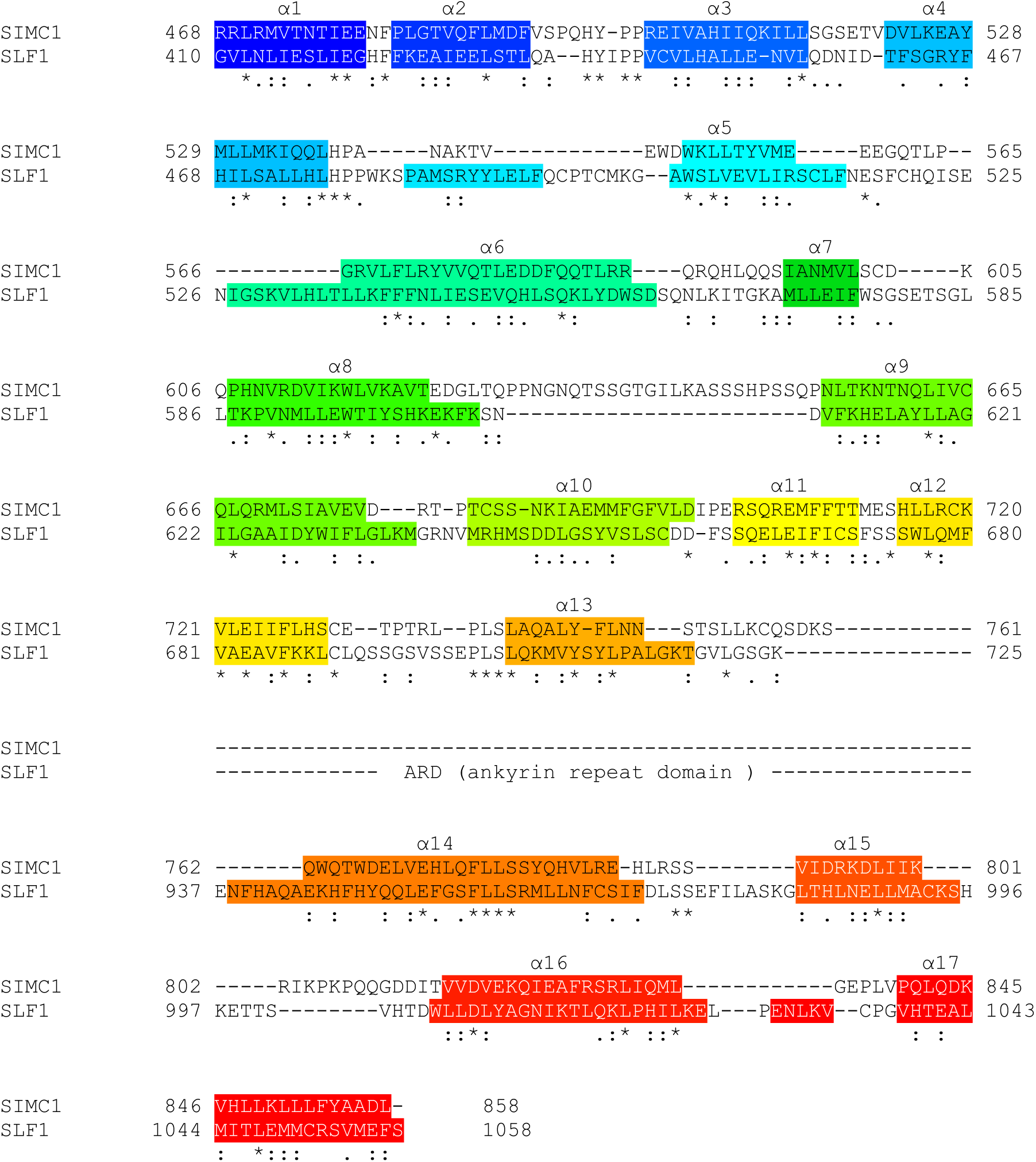
Structure-based sequence alignment of SIMC1 and SLF1. Secondary structures (α-helices) are colored.

**Figure 7 – figure supplement 5.**
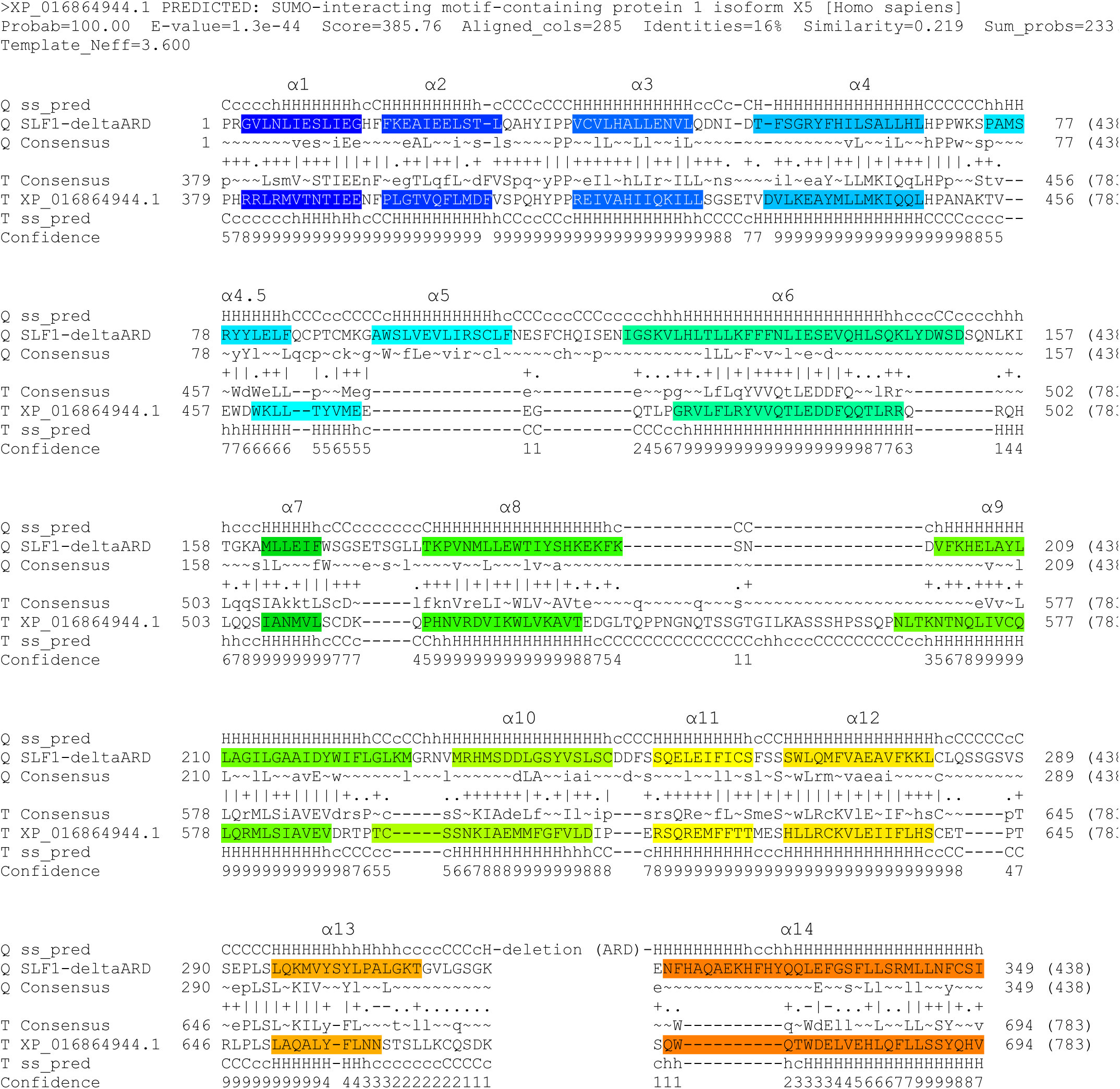
HHpred-aligned SLF1 and SIMC1 sequences. The query was SLF1^CTD_ΔARD^ (410-725 aa + 937-1058 aa), and SIMC1 was the top hit. The α-helices determined by AlphaFold (for SLF1) and cryo-EM (SIMC1) are colored.

## List of Supplementary files

Supplementary file 1 - Mass spectrometry identification of SMC5 interacting proteins labelled by SILAC and BioID.

Supplementary file 2 - Mass spectrometry identification of SIMC1 BioID. Supplementary file 3 - Key resources.

## List of source data

Figure 2 – source data 1. Full and unedited blots corresponding to panel C.

Figure 2 – figure supplement 2 – source data 1. Full and unedited blots.

Figure 2 – figure supplement 3 – source data 1. Full and unedited gels.

Figure 3 – source data 1. Full and unedited blots corresponding to panel D.

Figure 4 – source data 1. Primary data for graphs in panel A.

Figure 4 – source data 2. Primary data for graphs in panel C.

Figure 4 – source data 3. Full and unedited blots corresponding to panel D.

Figure 4 – figure supplement 1 – source data 1. Primary data for graphs in panel B.

Figure 4 – figure supplement 1 – source data 2. Full and unedited blots corresponding to panel C.

Figure 6 – source data 1. Full and unedited blots corresponding to panel B.

Figure 6 – source data 2. Primary data for graphs in panel D.

Figure 7 – source data 1. Full and unedited blots corresponding to panel D.

Figure 7 – source data 2. Full and unedited blots corresponding to panel E.

